# ER – lysosome contacts at a pre-axonal region regulate axonal lysosome availability

**DOI:** 10.1101/2020.06.16.153734

**Authors:** Nazmiye Özkan, Max Koppers, Inge van Soest, Alexandra van Harten, Nalan Liv, Judith Klumperman, Casper C. Hoogenraad, Ginny G. Farías

## Abstract

Neuronal function relies on careful coordination of organelle organization and transport. Kinesin-1 mediates transport of the ER and lysosomes into the axon and it is increasingly recognized that contacts between the ER and lysosomes influence organelle organization. However, it is unclear how organelle organization, inter-organelle communication and transport are linked and how this contributes to local organelle availability in neurons. Here, we show that somatic ER tubules are required for proper lysosome transport into the axon. Somatic ER tubule disruption causes accumulation of enlarged and less motile lysosomes at the soma. ER tubules regulate lysosome size and axonal translocation by promoting lysosome homo-fission. ER tubule – lysosome contacts often occur at a somatic pre-axonal region, where the kinesin-1-binding ER-protein P180 binds microtubules to promote kinesin-1-powered lysosome fission and subsequent axonal translocation. We propose that ER tubule – lysosome contacts at a pre-axonal region finely orchestrate axonal lysosome availability for proper neuronal function.

## INTRODUCTION

Neuronal organelle organization, functioning and transport must be carefully orchestrated to maintain neuronal architecture and function (Bentley and Banker, 2016; van Bergeijk et al., 2016). Microtubule (MT)-driven motor – organelle coupling ensures proper organelle transport into the two morphologically and functionally distinct structures of a neuron, the somatodendritic and axonal domains (Bentley and Banker, 2016; Britt et al., 2016; Gumy and Hoogenraad, 2018). From extensive studies in non-neuronal cells, it has been increasingly recognized that organelles form contacts with each other to execute essential processes such as lipid and ion transfer, organelle division and motor transfer (Raiborg et al., 2015b; Bonifacino and Neefjes, 2017; Wu et al., 2018). However, little is known about how organelle organization, inter-organelle communication and transport are linked and how this impacts local organelle availability in neurons.

The endoplasmic reticulum (ER) is one of the largest organelles and forms extensive contacts with various other organelles, including late endosomes (LEs)/ lysosomes (Friedman et al., 2013; Guo et al., 2018). The ER is organized as perinuclear ER cisternae connected with a network of ER tubules that spread into the cell periphery of unpolarized cells (Westrate et al., 2015; Zhang and Hu, 2016). In neurons, ER tubules are distributed along the somatodendritic and axonal domains, while ER cisternae are restricted to the somatodendritic domain (Wu et al., 2017; Farías et al., 2019). The shape of the ER is maintained by ER-shaping proteins such as reticulons (RTNs) and DP1, which induce the curvature of tubules, and CLIMP63, which generates flattened ER cisternae (Voeltz et al., 2006; Shibata et al., 2010; Westrate et al., 2015). Recent evidence has revealed that the ER is highly dynamic, undergoing fast remodeling in the order of seconds (Nixon-Abell et al., 2016; Guo et al., 2018). Although contacts between ER tubules and LEs/ lysosomes have been visualized in both unpolarized cells and in neurons from brain tissue (Friedman et al., 2013; Wu et al. 2017), it is less clear how ER remodeling regulates these organelle interactions.

It is well known that the ER and LEs/ lysosomes form contacts at membrane contact sites, where small molecules and lipids can be transported reciprocally (Wu et al., 2018; Lee and Blackstone, 2020). To maintain a steady state number and size and correct positioning of LEs/ lysosomes, essential for cellular homeostasis, they undergo series of fusion, fission and motor-based transport events (Saffi and Botelho, 2019). LE/ lysosome fission and motor loading onto LEs/ lysosomes often occurs in association to both the ER and MTs (Friedman et al., 2013; Rowland et al., 2014; Raiborg et al., 2015a; Guo et al., 2018). Yet, it remains unclear how local ER organization regulates LE/ lysosome size and how this is linked to motor transfer and MT interaction at contact sites.

Proper organization and transport of ER tubules and LEs/ lysosomes are crucial for neuronal development and function. ER tubules and LEs/ lysosomes are translocated from the soma into the axon by the kinesin-1 motor (Farías et al., 2017; Farías et al., 2019). Local availability of ER tubules instructs axon formation and regulates axonal synaptic vesicle cycling (Farías et al., 2019; Lindhout et al., 2019) and active transport of LEs/ lysosomes into the axon is required for proper clearance of faulty proteins and organelles located far away from the cell soma (Farías et al., 2017; Farfel-Becker et al., 2019). Interestingly, mutations in genes encoding ER-shaping proteins cause the neurodegenerative disease hereditary spastic paraplegia, in which aberrant lysosomes have been observed (Westrate et al., 2015; Allison et al., 2017; Lee and Blackstone, 2020). Therefore, it is important to understand how the organization of the ER and inter-organelle communication contribute to lysosome organization and local availability in neurons.

Here, we show that ER shape regulates local lysosome availability in neurons, in which somatic ER tubules promote lysosome translocation into the axon. Disruption of somatic ER tubules causes accumulation of enlarged and less motile mature lysosomes in the soma due to impaired lysosome homo-fission. We find that ER tubule – lysosome contacts are enriched in a pre-axonal region. The MT- and kinesin-1-binding ER protein P180 is enriched and co-distributed with kinesin-1-decorated axonal MT tracks in the same pre-axonal region, where it promotes lysosome motility, fission and axonal translocation. Together, our results support a model in which ER – lysosome contacts at a pre-axonal region finely orchestrate axonal lysosome availability.

## RESULTS

### ER shape regulates lysosome availability in the axon

To study the role of the ER in neuronal lysosome organization, we first investigated whether ER shape regulates lysosome distribution in primary cultures of rat hippocampal neurons. ER tubules are generated by two main ER tubule-shaping proteins, RTN4 and DP1, while flattened ER cisternae are maintained by CLIMP63. The abundance of these ER-shaping proteins regulates the conversion between cisternae and tubules (Voeltz et al., 2006; Shibata et al., 2010). We knocked down both RTN4 and DP1, or CLIMP63 and analyzed the distribution of GFP-tagged LAMP1, a marker for late endosomes and lysosomes (henceforth referred to as immature and mature lysosomes, respectively, or just lysosomes), in neurons at day-*in-vitro* 7 (DIV7). In control conditions, LAMP1-positive lysosomes were abundant in the soma and evenly distributed along dendrites and the axon (Figure 1A), as previously reported (Farías et al., 2017). Knockdown of RTN4 plus DP1 caused a drastic reduction of LAMP1-positive lysosomes along the axon, whereas CLIMP63 knockdown increased their axonal distribution (Figure 1A). Quantification of the polarity index (PI: [intensity dendrite – intensity axon] / [intensity dendrite + intensity axon]) confirmed the unpolarized distribution of lysosomes in the control condition (PI: 0.04). Removal of ER tubule-shaping proteins disrupted axonal lysosome distribution (PI: 0.4) whereas CLIMP63 knockdown neurons showed an increased axonal lysosome distribution (PI: −0.5) (Figure 1B). Similar results were observed using endogenously labeled LAMTOR4, another marker for lysosomes, in which ER tubule disruption caused an impaired LAMTOR4 distribution along the axon (Figures 1C and 1D).

**Figure 1.**
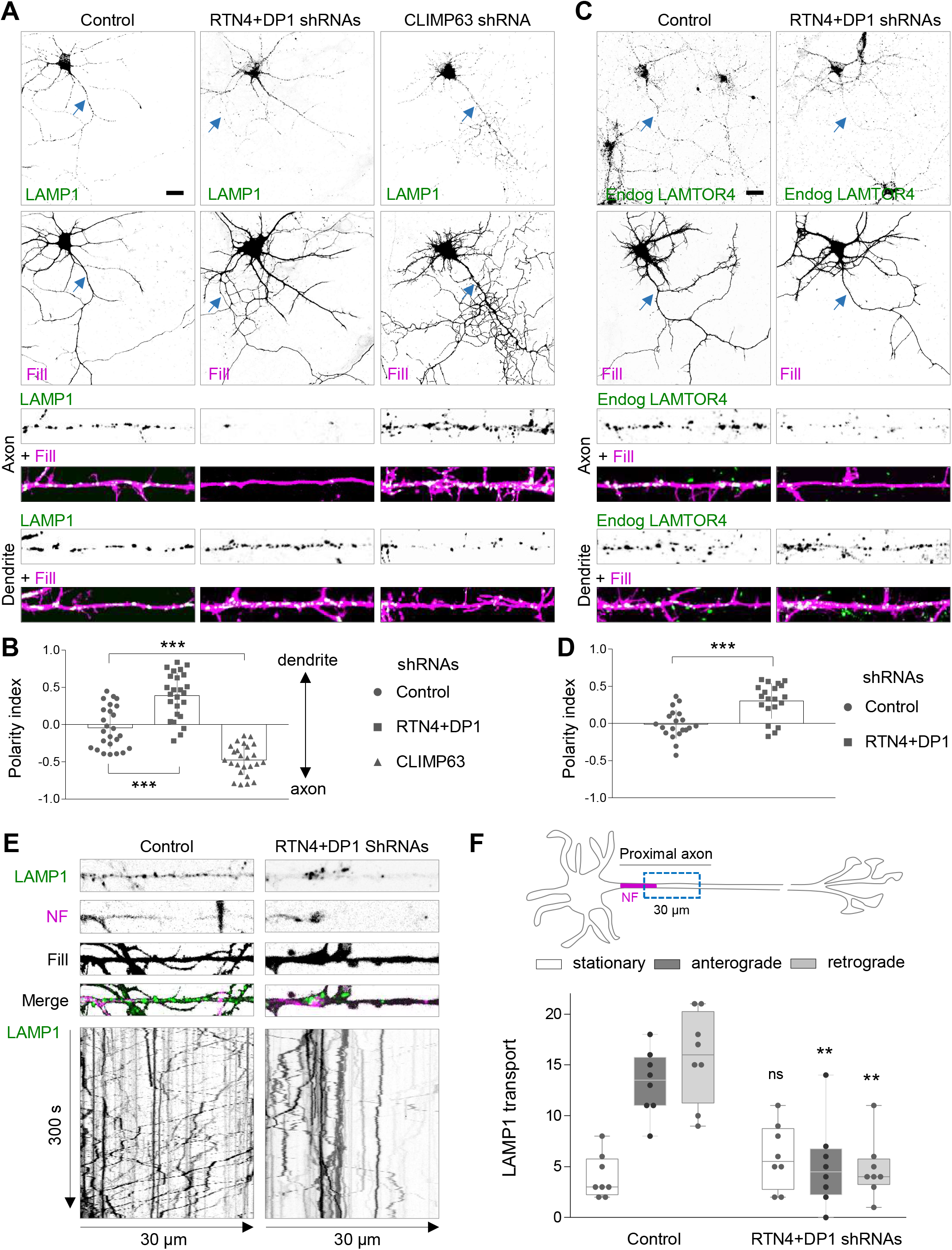
ER morphology controls lysosome translocation into the axon. (**A-B**) Representative images of DIV7 hippocampal neurons co-transfected at DIV3 with LAMP1-GFP (green), a mCherry fill (magenta) and a control pSuper plasmid or a pSuper plasmid containing a shRNA sequence targeting RTN4 plus DP1, or CLIMP63, in (A). Higher magnification of 40-μm straightened axon (top) or dendrite (bottom) segments. Quantification of LAMP1 polarity indices in (B). **(C-D)** Representative images of DIV7 neurons that were transfected with a mCherry fill (magenta) and a control pSuper plasmid or with shRNAs targeting RTN4 plus DP1 and stained with a LAMTOR4 antibody (green) in (C). Higher magnification of 40-μm straightened axon (top) or dendrite (bottom) segments. Polarity indices for LAMTOR4 in (D). **(E)** Representative still images (top) and kymographs (bottom) from a 30-μm-segment of straightened proximal axons of live neurons co-transfected with LAMP1-GFP (green) and fill (gray) together with control pSuper or shRNAs targeting RTN4 plus DP1, and labelled for the axon initial segment (AIS) marker Neurofascin (NF; magenta) prior to imaging for 300 s. See also Video S1 **(F)** Quantification of LAMP1-positive lysosome movement. Schematic representation of a neuron indicating the axonal region selected for quantification (top) and average number of stationary, anterograde and retrograde pools (bottom). Blue arrows point to the proximal axon in (A) and (C). Scale bars represent 20 μm in (A) and (C). Boxplots show the mean and individual datapoints each represent a neuron, in (B), (D), and (F); ns-not significant, ***p<0.001 and **p<0.01 comparing conditions to control (Kruskal-Wallis test followed by a Dunn’s multiple comparison test) in (B) and (F), and (*t*-test) in (D).

Reduced distribution of lysosomes along the axon could be explained by an increased retrograde transport of lysosomes from the axon into the soma or an impaired translocation of lysosomes from the soma into the axon. To study this, LAMP1 dynamics was analyzed by live-cell imaging in a 30-μm-length segment of the proximal axon during a period of 300 seconds (Figures 1E and 1F). In control neurons, an average of 29 out of 33 LAMP1-positive lysosomes per neuron were motile at the proximal axon, from which 13 transported anterogradely into the axon tip and 16 transported retrogradely to the cell soma (Figure 1F; Video S1). Knockdown of RTN4 plus DP1 caused a reduction of 51.5% in the total number of lysosomes distributed along the proximal axon, decreasing both antero- and retrograde LAMP1 movement, while the stationary pool remained unaffected (Figures 1E and 1F; Video S1). These results show that ER tubules play a critical role in regulating lysosome translocation from the soma into the axon.

### Somatic, but not axonal, ER tubules promote lysosome translocation into the axon

Since most of the axonal ER corresponds to ER tubules (Wu et al., 2017; Farias et al., 2019), we wondered whether the contact between ER tubules and lysosomes locally regulates lysosome distribution and dynamics (Figure 2A). To study this, we used a heterodimerization system to control ER tubule positioning. We induced a sustained retention of ER tubules in the somatodendritic domain, by triggering the binding of the KIFC1 (a minus-end driven motor) to ER tubules by fusing a Streptavidin (Strep) sequence to KIFC1 and a SBP to GFP-tagged RTN4A (Figure 2B). This strategy allows local axonal depletion of ER membranes, as previously confirmed by the absence of several other ER markers in the axon (Farías et al., 2019). Lysosome distribution in neurons was analyzed after 24-48 hours of co-expression of LAMP1-RFP and GFP-SBP-RTN4 in the presence or absence of KIFC1-Strep. In the control condition, LAMP1 and SBP-RTN4 were co-distributed along the entire neuron, in both the somatodendritic and axonal domains (Figure 2A). In the presence of KIFC1-Strep, axonal ER tubules containing SBP-RTN4 were pulled from the axon into the somatodendritic domain, while lysosomes were still distributed along the axon (Figure 2B). We then analyzed whether axonal ER tubule removal affected the dynamics of lysosomes along the axon. Co-expression of SBP-RTN4 and KIFC1-Strep, did not cause a reduction in axonal LAMP1 motility. Antero- and retrograde movement as well as the stationary pool of LAMP1-positive lysosomes were similar to control neurons expressing only SBP-RTN4 (Figures 2D and 2E, Video S2). These results indicate that axonally distributed ER tubules do not contribute to the availability and dynamics of lysosomes along the axon. To further examine the role of somatic ER tubules, we fused a Strep sequence to the axonal plus-end driven kinesin-1 motor KIF5A (Figure 2C). Co-expression of SBP-RTN4A and Strep-KIF5A induced axonal transport of ER tubules and their accumulation in the distal axon (Figure 2C). In these neurons, LAMP1-positive lysosomes were distributed in the somatodendritic domain but drastically reduced along the axon (Figure 2C). Live-cell imaging showed that the total number of lysosomes along the proximal axon was impaired, and the bidirectional movement of lysosomes was drastically reduced compared to control neurons, while the stationary pool remained unaffected (Figures 2D and 2E; Video S2). These results indicate a role for somatic ER tubules in promoting lysosome translocation into the axon.

**Figure 2.**
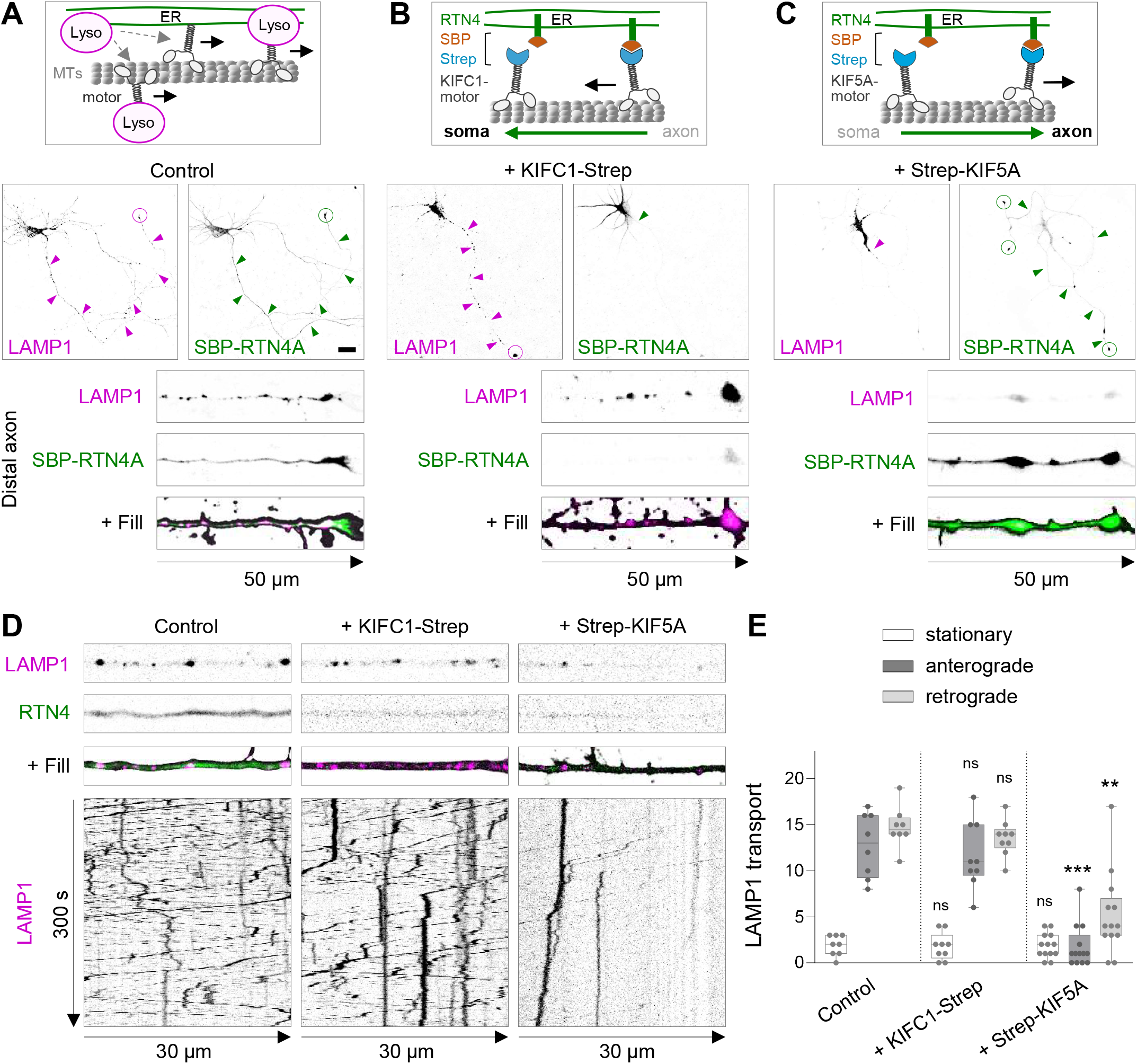
Somatic ER tubules control lysosome translocation into the axon. **(A)** Schematic model for motor-driven lysosome transport regulated by ER-tubules (top). Representative images of LAMP1-RFP (magenta) and GFP-SBP-RTN4 (green) distribution in a control DIV6 neuron co-transfected with fill (middle), and higher magnification of distal axon (bottom). **(B)** Schematic representation of Streptavidin (Strep)-SBP heterodimerization system using SBP-RTN4 and Strep-KIFC1 for MT-dependent minus-end ER-tubule transport and its persistent somatic retention (top). Representative images of LAMP1-RFP (magenta) and GFP-SBP-RTN4 (green) distribution in a neuron co-transfected with Strep-KIFC1 and fill (middle) and higher magnification of distal axon (bottom). **(C)** Schematic representation of Strep-SBP system using SBP-RTN4 and KIF5A-Strep for MT-dependent anterograde transport of ER-tubules and its persistent distribution in distal axons (top). Representative images of LAMP1-RFP (magenta) and GFP-SBP-RTN4 (green) distribution in a neuron co-transfected with KIF5A-Strep and fill (middle) and higher magnification of distal axon (bottom). **(D-E)** Representative still images (top) and kymographs (bottom) from a proximal axon of live neurons co-transfected with LAMP1-RFP (magenta) and SBP-RTN4 (green), in absence of a motor protein (control), or with Strep-KIFC1 or KIF5A-Strep (from left to right) in (D). Quantification of stationary, anterograde and retrograde movement of lysosomes from conditions in D, in (E). See also Video S2. Magenta and green arrows point to the abundance of LAMP1 and SBP-RTN4 along the axon and dashed circles point to their accumulation at axon tips. Scale bars represent 20 μm in (A), (B) and (C). Boxplot shows the mean and individual datapoints each represent a neuron; ns-not significant, ***p<0.001 and **p<0.01 comparing conditions to control (Kruskal-Wallis test followed by a Dunn’s multiple comparison test) in (E).

### Local ER tubule disruption causes enlarged and less motile mature lysosomes in the soma

To determine how disruption of ER tubules impairs lysosome translocation into the axon, we analyzed the organization of lysosomes in the soma by confocal imagining and z-stack reconstruction. This revealed that ER tubule disruption caused a striking enlargement of LAMP1- or LAMTOR4-positive lysosomes compared to control neurons (Figures 3A and 3B). We found that these enlarged lysosomes were often less motile (Video S3). Similarly, somatic ER tubule redistribution into the axon induced an enlargement of lysosomes in the soma (Figure 3C). These enlarged lysosomes were also less dynamic and unable to translocate into the axon and were retained at a region preceding the axon initial segment (Videos S4 and S5). Conversely, redistribution of ER tubule into the soma caused a decreased lysosome size together with an increased motility of lysosomes compared to control neurons (Video S4).

**Figure 3.**
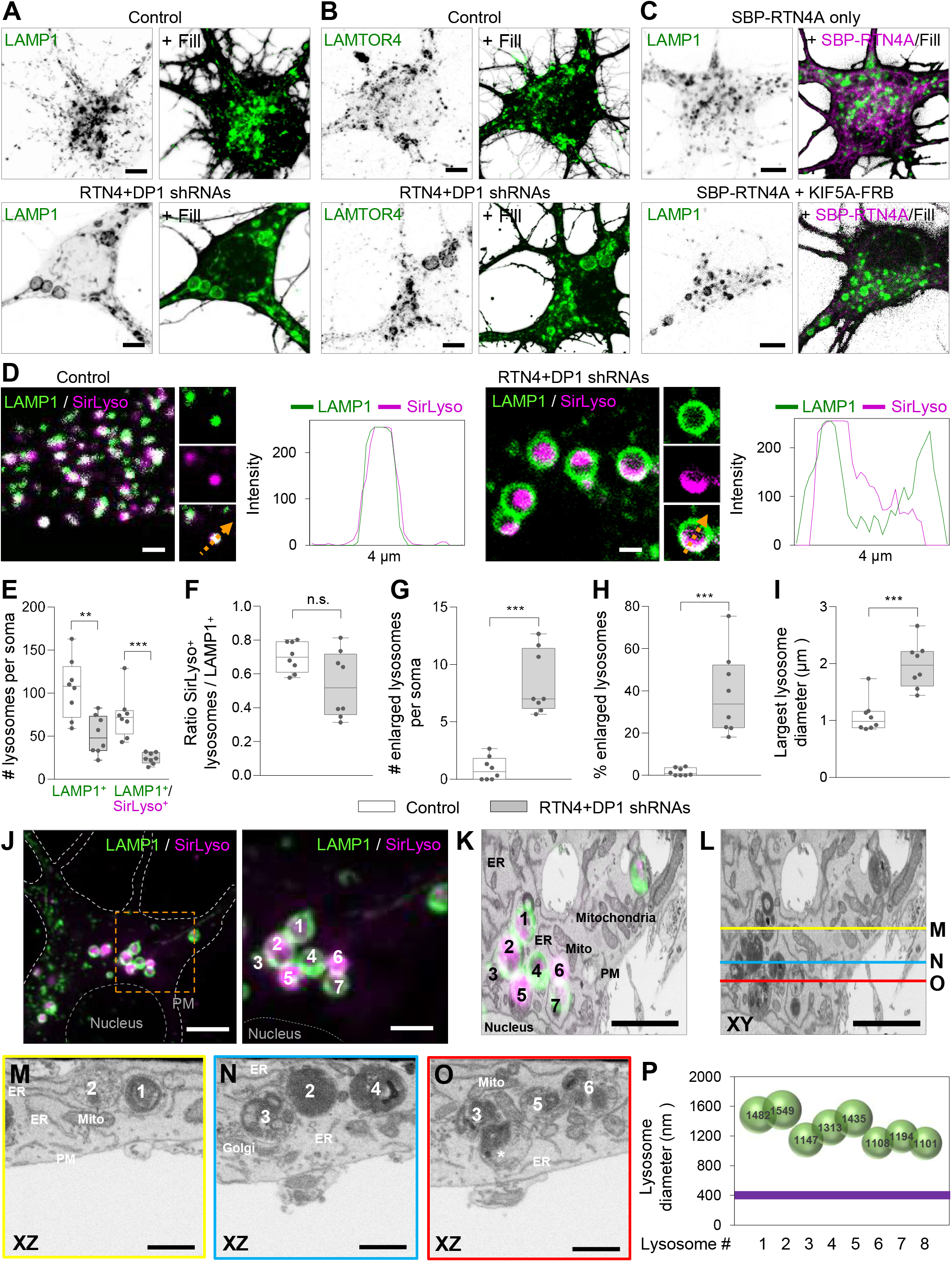
ER tubule disruption causes enlargement and reduced motility of mature lysosomes. **(A-B)** Representative images of lysosomes distributed in the soma of DIV7 neurons transfected at DIV3 with fill and a control pSuper plasmid (top) or pSuper plasmids containing shRNAs targeting RTN4 plus DP1 (bottom) together with LAMP1-GFP (A) or stained for endogenous LAMTOR4 (B), green in merges. **(C)** Representative images of lysosomes in the soma of DIV6 neurons co-transfected at DIV5 with LAMP1 (green) and fill, together with only SBP-RTN4 (magenta) as a control (top) or SBP-RTN4 plus KIF5A-Strep (bottom) to pull ER-tubules into the axon. See also Video S3, S4 and S5. **(D)** Representative still images of the soma of DIV7 neurons transfected as in (A) and labelled live for active cathepsin-D (magenta) with SirLyso. LAMP1 in green. The size of LAMP1-positive lysosomes and luminal distribution of cathepsin content is shown. Intensity profile line on the right of magnified image of a lysosome for each condition. See also Figures S1A-E and Video S6. **(E-I)** Parameters indicated in each graph were quantified from the soma of neurons transfected and labeled as in (D). Control, white bars; RTN4 plus DP1 knockdown, gray bars. **(J-P)** Correlative light electron microscopy (CLEM) of enlarged lysosomes. (J) FM image of a fixed neuron, knock-down for RTN4 and DP1 and expressing LAMP1-GFP. SirLyso indicates hydrolase active lysosomes. Nucleus and plasma membrane (PM) are indicated with dashed lines. A cluster of enlarged lysosomes was selected (orange rectangle representing the ROI) for 3D-EM analyses. The right panel shows an enlargement of the ROI with 7 marked lysosomes. (K) Reconstructed FIB.SEM slice of ROI in same (XY) orientation as FM and with overlay of FM signal. (L) Same EM image as in (K) marked with yellow, blue, and red lines that correspond to the orthogonal images shown below. (M) XZ plane image corresponding to the yellow line in (L) and showing cross sections of lysosomes #1 and #2. Lysosome #2 shows many intraluminal vesicles in this plane corresponding to the SirLyso signal in the FM image. (N) XZ plane image corresponding to the blue line in (L) and showing cross sections of Lysosomes #2 and #4. In contrast to (M), lysosome #2 contains dense degraded material in this plane, showing the compartmentalized content of these enlarged lysosomes. Lysosome #3 and #4 are closely interacting. (O) XZ plane image corresponding to the red line in (L) showing lysosomes #3, #5 and #6. An additional lysosome (*) tightly in contact with lysosome #3 is seen in EM but not visible in the FM image. Many interactions between lysosomes were observed, but they remained separate entities. PM, Golgi, ER and mitochondria present in the EM ROI, are indicated in (K), (M), (N), and (O). (P) Size distribution plot of the 8 lysosomes visible in the EM images. All lysosomes are remarkably similar in size and shape, with a globular form of 1100 – 1500 nm diameter, 3 times bigger than the average lysosome size (400 nm; purple line). See also Videos S7 and S8. Scale bars represent 5 μm in (A), (B), (C), (J), 2 μm in (D), (J, right panel), (K) and (L), and 1 μm in (M), (N) and (O). Boxplots show the mean and individual datapoints each represent a neuron; ns-not significant, ***p<0.001 and **p<0.01 comparing conditions to control (Mann-Whitney U) in (E), (F), (G), (H) and (I).

Next, we analyzed whether disruption of ER tubules alters the maturation state of these enlarged lysosomes, by analyzing the presence of active cathepsins in LAMP1-positive lysosomes (mature lysosomes). We tested two probes in live neurons; Magic-Red, which becomes fluorescent after cathepsin-B breaks down the substrate, and SirLyso, which labels active cathepsin-D (Ferguson, 2018). We found that ER tubule disruption did not affect lysosome activity, as most of the less motile enlarged lysosomes contained active cathepsins, and this lysosome activity was often observed compartmentalized within their luminal domain (Figure 3D; Figures S1A and S1B; Video S6). We quantified the total number of lysosomes and mature lysosomes per soma in live neurons. The total number of LAMP1-positive lysosomes was reduced to 49% after ER tubule knockdown compared to control neurons, from which the mature lysosome population (LAMP1 / SirLyso positive) was reduced to 33% (Figure 3E). The proportion of mature lysosomes to all LAMP1-positive lysosomes was not significantly reduced after disruption of ER tubules (Figure 3F). Quantification of the number of lysosomes by size revealed an average of 8.4 mature lysosomes per soma with a diameter bigger than 1 μm (considered as enlarged; de Araujo et al., 2020) after ER tubule knockdown compared to only 0.9 large mature lysosomes per soma in control neurons (Figure 3G). The percentage of large lysosomes relative to all LAMP1-positive lysosomes, revealed that around 1.9% of mature lysosomes were larger than 1 μm in control neurons, while this number increased to 39% after ER tubule disruption (Figure 3H). The average diameter of the largest mature lysosome per soma was doubled compared to control neurons (Figure 3I).

To reveal the ultrastructural morphology of the enlarged LAMP1-positive lysosomes we performed correlative light electron microscopy (CLEM), by which we selected a cluster of LAMP1 and SirLyso-positive organelles for FIB.SEM (focused ion beam scanning electron microscopy) imaging (Fermie et al., 2018) (Figures 3J-O; Videos S7 and S8). 3D analysis showed that the enlarged lysosomes have a consistent globular shape of 1100 – 1500 nm diameter. For comparison, control lysosomes are on average 400 nm in size and more variable in shape (de Araujo et al., 2020) (Figure 3P). The content of the aberrant lysosomes was a heterogenous mix of dense, degraded material and accumulations of intraluminal vesicles (Figures 3L-O). The compartmentalized fluorescence SirLyso signal corresponded to the areas with intraluminal vesicles, indicating that these membranes are subject to lysosomal degradation. The lysosomes showed many interaction sites with each other which extended over considerable distances, but they remained clearly separate entities. These data show that disruption of ER tubules leads to the accumulation of a collection of enlarged, enzymatically active lysosomes of remarkable consistent size and shape.

Altogether, these results show that somatic ER tubules control axonal lysosome availability by regulating lysosome size and motility but not activity.

### ER tubules regulate lysosome homo-fission

Lysosome size is carefully controlled by balancing hetero- or homo-fusion and fission events (Saffi and Botelho, 2019). The enlarged LAMP1-positive structures we observed could therefore be caused by increased fusion and/or reduced fission. We first analyzed whether enlarged lysosomes were fused with other components of the endo-lysosomal system, including early endosomes, recycling endosomes and autophagosomes, labeled by endogenous EEA1, GFP-tagged Rab11 and endogenous p62, respectively. The enlarged LAMP1- or LAMTOR4-positive lysosomes caused by ER tubule disruption were not particularly enriched for EEA1, Rab11 or p62 markers (Figures S1C-E). This suggests that the enlargement of lysosomes is not induced by a mechanism that involves an increase in hetero-fusion or reduction in hetero-fission between lysosomes and early endosomes, recycling endosomes or autophagosomes.

Then, we examined whether an increase in homo-fusion and/or a reduction in homo-fission events could explain our observed enlarged lysosomes. To determine whether homo-fusion and homo-fission were altered by ER tubule disruption, neurons expressing LAMP1-GFP were labelled for SirLyso and imaged in the soma for a period of 300 seconds (Video S6). We focused on fusion and fission events occurring with mature lysosomes since these were more affected by ER tubule disruption. In control neurons, we observed fusion events between mature lysosomes positive for both LAMP1-GFP and SirLyso with immature lysosomes positive only for LAMP1-GFP, as well as fusion between mature lysosomes (Figures 4A and 4B; Video S9). Fission events were also often observed, including budding of an immature or mature lysosome from a spherical mature lysosome, as well as budding from the tubular domain of a tubular-shaped mature lysosome (Figures 4C-E; Video S9). In neurons with disrupted ER tubules, fusion events between mature and immature lysosomes and between mature lysosomes were also observed (Figures 4F and 4G; Video S9). ER tubule disruption clearly affected lysosome fission. Enlarged lysosomes often failed in the termination of the budding process to generate a mature or immature lysosome from a parent mature lysosome (Figures 4H and 4I; Video S9). In addition, enlarged lysosomes generated instable tubules undergoing elongation followed by retraction after unsuccessful tubule fission (Figure 4J; Video S9). Quantification of the number of fusion events per soma showed that an average of 135 fusion events occurred in the soma of control neurons during a period of 300 seconds. We unexpectedly found that ER tubule disruption caused a reduction in fusion events per soma from an average of 135 events in control neurons to 30 events in ER tubule knockdown neurons (Figure 4K).

**Figure 4.**
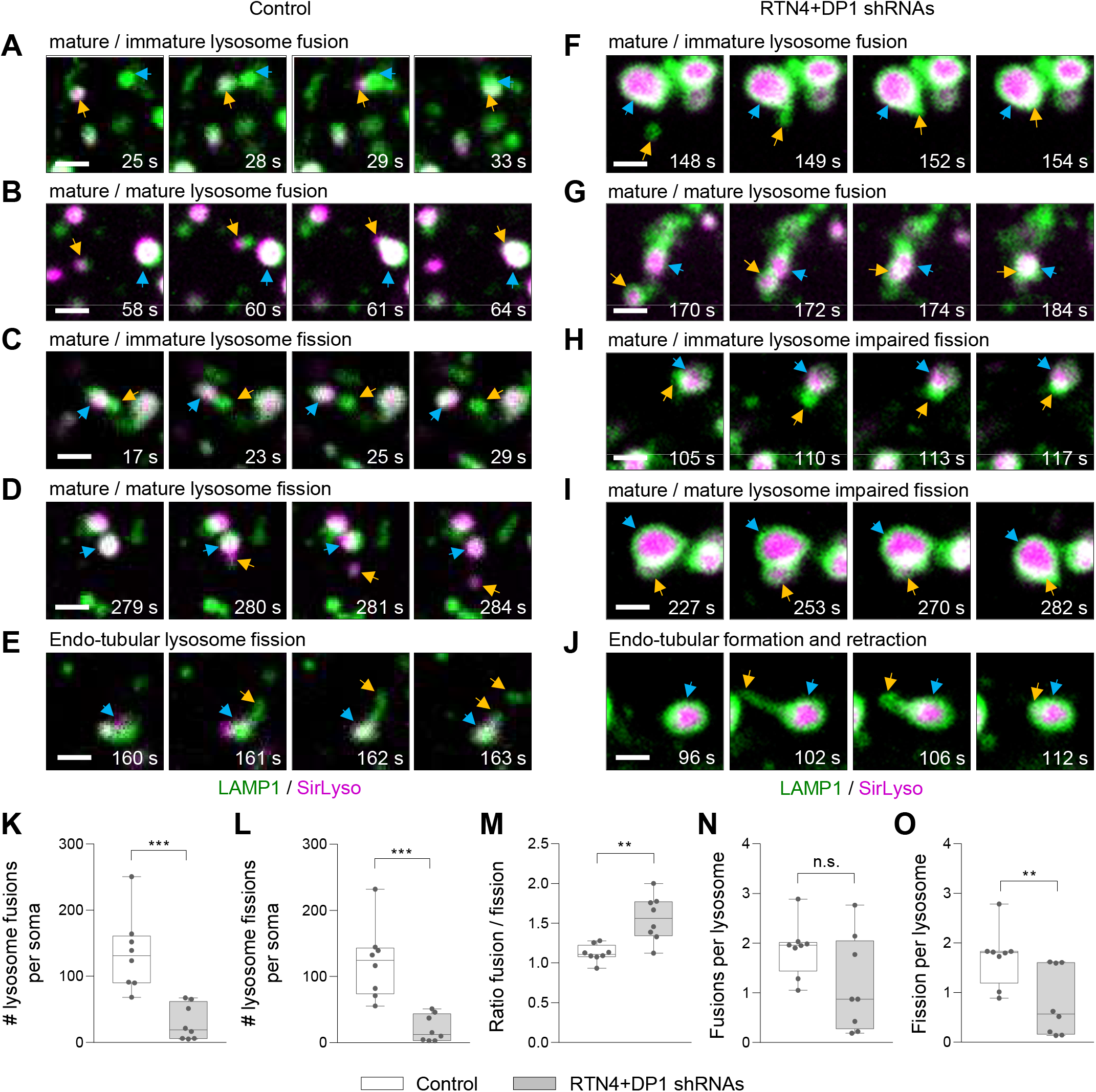
Enlarged lysosomes are caused by an imbalance in homo-fission. **(A-J)** Representative still images of fusion and fission events from DIV7 neurons co-transfected with control pSuper (A-E) or shRNAs targeting RTN4 plus DP1 (F-J) together with LAMP1-GFP (green) and labelled with SirLyso (magenta) prior imaging for 300 s every 1 s. Fusion between lysosomes in (A), (B), F), (G); Fission in (C), (D), E); impaired fission in (H), (I), (J). Time scale included per event. See also Video S9. Blue and orange arrows point to two lysosome undergoing fusion, or one lysosome budding from a parent lysosome. In all images, scale bars represent 1 μm. **(K-O)** Parameters indicated in each graph were quantified from live neurons transfected and labelled as in (A-J) and imaged for 300 s every 1 s. Control, white bars and RTN4 plus DP1 knockdown, gray bars. Boxplots show the mean and individual datapoints each represent a neuron; ns-not significant, ***p<0.001 and **p<0.01 comparing conditions to control (Mann-Whitney U).

Fission events were also reduced from an average of 122 events in control neurons to 21 events in neurons with ER tubule knockdown (Figure 4L). The number of fusion and fission events per soma were similar in control neurons with a fusion/fission ratio of 1.1, while ER tubule disruption caused an increase in this ratio to 1.6 (Figure 4M). Since RTN4/DP1 knockdown also reduced the total number of lysosomes (Figure 3E), we calculated the number of fusion and fission events per lysosome for each soma. This revealed that fusion events per lysosome were not significantly affected, while fission events were significantly reduced compared to control neurons (Figures 4N and 4O). These results are consistent with previous work in non-neuronal cells, in which ER – endosome contacts contribute mainly to endosome fission, but not fusion (Friedman et al., 2013; Rowland et al., 2014; Hoyer et al., 2018). Together, these results suggest that somatic ER tubule – lysosome contacts control lysosome fission to regulate lysosome size and translocation in neurons.

### ER tubule – lysosome contacts occur at the soma and are enriched at a pre-axonal region

We wanted to confirm that ER tubules regulate lysosome size and axonal translocation via a direct local contact between these two organelles in the soma. To visualize the distribution of ER – lysosome contacts in neurons, we utilized the proximity-based split-APEX labelling assay (Han et al., 2019). In this assay, a split version of APEX2, an engineered peroxidase able to covalently tag proximal endogenous proteins with biotin in living cells, is used. Two inactive fragments, AP and EX, can only reconstitute driven by a molecular interaction, resulting in biotinylation of a contact site (Figure 5A; Han et al., 2019). We tagged protrudin, an ER tubule protein enriched in contact sites (Raiborg et al., 2015a), with an AP module and a V5-tag, and the endosome and lysosome adaptor Rab7 with an EX module. Neurons expressing the split-APEX system showed a clear co-distribution of AP-protrudin and the endogenously labelled lysosome marker LAMTOR4 (Figure 5B). The biotinylation around these two organelles, detected by fluorescently labelled streptavidin (Strep-Alexa568), indicated their co-distribution corresponds to a true contact (Figures 5B and 5G). The Strep signal was specific, as only the incubation with hydrogen peroxide, which catalyzes the proximity labelling reaction, produced biotinylation (Figure 5G). Importantly, we observed that ER – lysosome contacts were formed mainly in the soma and they were particularly enriched in a pre-axonal region (Figures 5C and 5D). ER tubule disruption caused a dramatic reduction of 91% in streptavidin intensity, compared to the control condition (Figures 5E-G). This experiment also confirmed that these contacts occur mainly between ER tubules and lysosomes, as ER tubule disruption caused the redistribution of protrudin within more flattened ER cisternae that are excluded from the pre-axonal zone (Figure 5E), as previously reported (Farías et al., 2015). Enlarged lysosomes accumulated at the pre-axonal zone and were prevented from entering the proximal axon (Figure 5F). Thus, ER tubules and lysosomes form contacts in the soma which are enriched in a pre-axonal region, supporting a direct role for ER tubules in lysosome fission and subsequent translocation into the axon.

**Figure 5.**
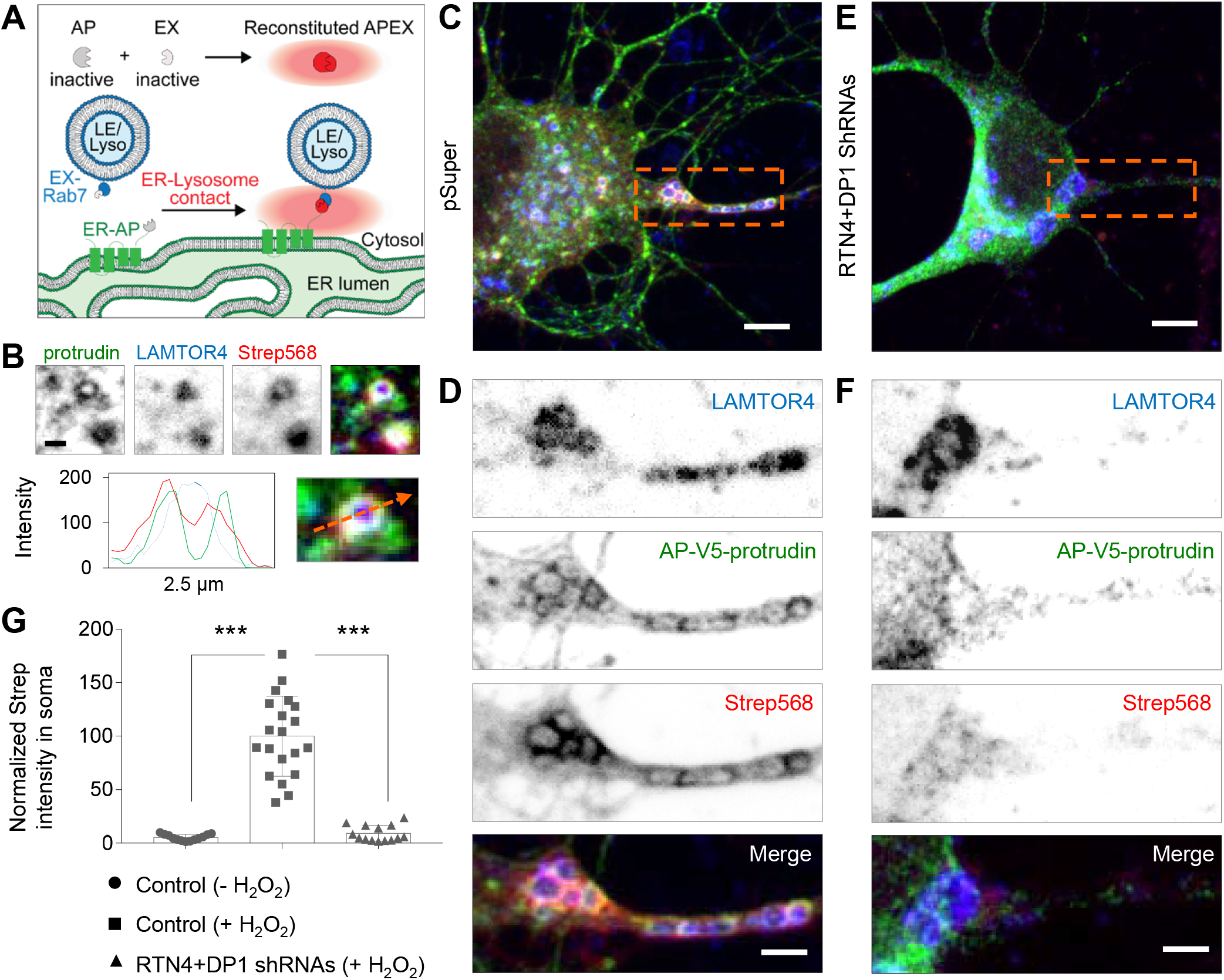
ER – lysosome contacts are enriched in a somatic pre-axonal region and they require ER tubule formation. **(A)** Schematic representation of Split-APEX system used to visualize ER – lysosome contacts. An ER tubule-contact marker (protrudin) is fused to an AP module and the lysosome adaptor Rab7 is fused to an EX module. Only proximity of the two proteins allows reconstitution of full APEX2. After addition of biotin-phenol (BP) and H2O2, APEX2 biotinylates proteins in close proximity and this can be detected by fluorescently conjugated-streptavidin (Strep; red radius). **(B)** Representative images of a magnified region from the soma of a neurons transfected with AP-V5-protrudin (green) and EX-Rab7, treated with biotin-phenol (BP) and H2O2, and labelled with an antibody against LAMTOR4 (blue) and Alexa568-conjugated Strep (red). Intensity profile line, bottom. **(C-F)** Representative images of neurons transfected, treated, and labelled as in (B), plus control pSuper plasmid (C) and (D) or shRNAs targeting RTN4 plus DP1 (E) and (F). Higher magnification of dashed orange boxes in (C) and (E) are shown in (D) and (F). **(G)** Average Strep intensity in soma of neurons transfected as in (C-F). Control neurons treated only with BP or control and knockdown neurons treated with BP plus H2O2 were labelled as in (B). Scale bars represent 5 μm in (D) and (F), 2 μm in (E), (G), and 1 μm in (B). Boxplot shows the mean and individual datapoints each represent a neuron; ***p<0.001 comparing conditions to control (ANOVA test followed by a Tukey’s multiple comparisons test) in (G).

### P180, a kinesin-1-binding protein enriched in ER tubules at the pre-axonal region, promotes lysosome translocation into the axon

The kinesin-1 motor has been shown to preferentially bind axonal microtubules in the soma in a region preceding the axon initial segment to promote organelle translocation into the axon (Farías et al., 2015; Farías et al., 2017; Farías et al., 2019). Besides its role in organelle transport, kinesin-1 has also been shown to promote organelle fission by generating the forces required for organelle budding (Du et al., 2016). Since our findings support a model in which ER tubules contact lysosomes at the pre-axonal region to regulate lysosome size for proper lysosome translocation into the axon, we searched for an ER protein that could mediate this process. This protein should be enriched at the pre-axonal region and should be able to bind kinesin-1. Three ER proteins containing a kinesin-1-binding domain, protrudin, KTN1, and P180, have previously been proposed to act as membrane anchor proteins that couple organelles to kinesin-1 for organelle translocation (Raiborg et al., 2015a; Matsuzaki et al., 2011; Ong et al., 2020; Diefenbach et al., 2004). To study whether kinesin-1-binding ER proteins are involved in lysosome translocation into the axon, we knocked down protrudin, KTN1 and P180 in neurons and analyzed lysosome distribution. We observed that only P180 knockdown reduced lysosome distribution in the axon (Figures S2A and S2B). P180 knockdown reduced lysosome translocation into the axon, as the total number of moving lysosomes was decreased, while stationary lysosomes remained unaffected in the proximal axon; a phenotype similar to what we observed after ER tubule disruption (Figures 6A and 6B, Figures 1E and 1F). Knockdown of P180 also resulted in enlarged lysosomes that accumulated in a pre-axonal region (Figure 6C; Video S10).

**Figure 6.**
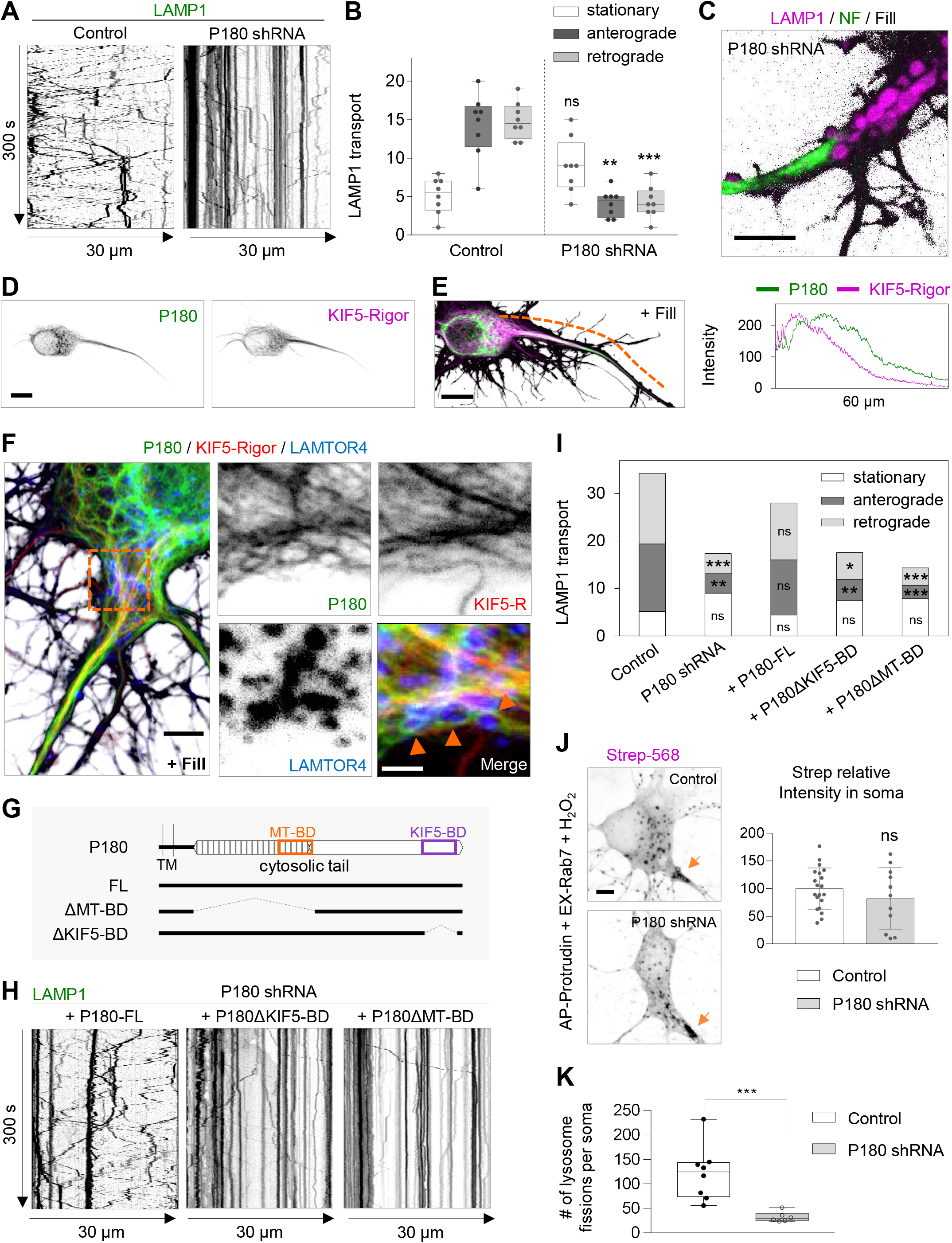
The KIF5-binding and ER protein P180 is enriched in a pre-axonal region and required for axonal lysosome translocation but not for ER – lysosome contact formation. **(A-C)** Representative kymographs of DIV7 neurons transfected with LAMP1-GFP and fill together with control pSuper plasmid or shRNAs targeting P180 in (A). Lysosome movement at the proximal axon was imaged for 300 s every 1 s. Quantification of lysosome movement in the proximal axon, in (B). Representative still image from Video S10 of the pre-axonal – AIS region of a neuron transfected with LAMP1-GFP (magenta), fill (gray) and stained for NF (green) in (C). See also Figures S2A and S2B. **(D-E)** Representative images of neurons transfected with mCherry-P180 (colored green), GFP-KIF5A-Rigor (colored red) and fill in (D). Merged image and intensity profile line from dashed orange segment, in (E). See also Figures S3A-C. **(F)** Representative image of a neuron transfected as in (D-E) and stained for LAMTOR4. Right panels show higher magnification of dashed line square. **(G)** Schematic representation of P180 protein with its short luminal domain, transmembrane domain (TM), microtubule binding domain (MT-BD in orange box) and KIF5-motor binding domain (KIF5-BD in purple box. Three constructs of P180 protein were generated as full length (FL), MT-BD deleted construct (ΔMT-BD) and KIF5-BD deleted construct (ΔKIF5-BD). **(H-I)** Representative kymographs of lysosome movement from neurons transfected as in (A) together with shRNA-resistant P180-FL, P180ΔKIF5-BD or P180ΔMT-BD constructs in (H). Control pSuper and shRNAs for P180 alone are shown in (A). Quantification of lysosome movement in the proximal axon in (I). **(J)** Representative images of neurons expressing split-APEX system and treated as in Figure 5B, co-expressing a control pSuper vector or shRNAs targeting P180 and labelled with Alexa568-conjugated Strep. Arrows point to a pre-axonal region. Graph shows the relative streptavidin intensity in control neurons versus shP180 treated neurons. n=20 and 11 neurons, respectively. **(K)** Quantification of lysosome fission events from live neurons transfected and labelled as in Figure 4 and imaged for 300 s every 1 s. Control, white bars (same as in Figure 4) and P180 knockdown, gray bars. Knockdown experiments performed in the same day than control neurons. Boxplots show the mean and individual datapoints each represent a neuron; ns-not significant and ***p<0.001 comparing conditions to control (Mann-Whitney U). See also Figures S2C-H and Video S11. Scale bars represent 10 μm in (D) and (E), 5 μm in (C), (F) and (J), and 2 μm in (F, right panels). Boxplot shows the mean (I) or mean and individual datapoints in (B) and (J); ns-not significant, ***p<0.001 and **p<0.01 comparing conditions to control (Kruskal-Wallis test followed by a Dunn’s multiple comparison test) in (B) and (I), and (Mann-Whitney U) in (J).

We previously showed that P180 is enriched in ER tubules preceding the axon initial segment (Farías et al., 2019). P180 and kinesin-1 KIF5A-Rigor, a motor mutant that can bind to but not walk or dissociate from microtubules, co-distributed in a pre-axonal region, indicating this is their main site of interaction (Figures 6D and 6E). Moreover, we observed lysosomes in contact with P180-enriched ER tubules in close proximity to kinesin-1-decorated microtubule tracks in this region (Figure 6F). The enrichment of P180 in a pre-axonal region required ER tubule formation, as ER tubule disruption caused the re-distribution of P180 into somatic ER cisternae absent from a pre-axonal region (Figure S3).

Besides a kinesin-1-binding domain, P180 contains a microtubule-binding domain located in a basic decapeptide repeat domain involved in ER tubule – MT co-stabilization (Ogawa-Goto et al., 2007; Farías et al., 2019). To determine whether the kinesin-1-binding domain (BD) and/or the MT-BD of P180 are required for axonal lysosome translocation, we performed knockdown and rescue experiments with shRNA-resistant P180 constructs (Figure 6G). Co-expression with full-length P180 rescued the reduced translocation of lysosomes into the axon after P180 knockdown, while co-expression with P180ΔKIF5-BD or P180ΔMT-BD deletion constructs was not sufficient to rescue this phenotype. This indicates that both domains are required for lysosome translocation into the axon (Figures 6H and 6I). Then, we investigated whether P180 is important for the formation of ER – lysosome contacts by using the split APEX assay (Figure 5A). However, knockdown of P180 did not reduce the streptavidin signal generated by AP-protrudin and EX-Rab7 (Figure 6J), indicating that P180 is not involved in ER tubule – lysosome contact formation (Figure 6C). Finally, we found that P180 knockdown reduced lysosome motility and often resulted in an agglomeration of mature lysosomes at a pre-axonal region (Video S11). We also observed a significant reduction in lysosome fusion and fission events, with no drastic reduction in the number of lysosomes (Figure 6K, Figures S2C-H). This is likely a consequence of the reduced motility that impairs the ability of lysosomes to complete fission and undergo new fusion events with other lysosomes (Video S11). Together, these results indicate that P180, enriched in ER tubules at a pre-axonal region, interacts with axonal microtubules, and is required for lysosome translocation into the axon. We find that P180 is not essential for ER – lysosome contact formation between ER and lysosomes. However, it may participate in contact – MT stabilization for subsequent kinesin-1-powered lysosome fission and translocation, as both its MT- and kinesin-1-binding domains are required for axonal lysosome translocation.

## DISCUSSION

Here we propose a model in which ER tubule – lysosome contacts at a pre-axonal region promote kinesin-1-powered lysosome fission and subsequent translocation into the axon. We show that ER shape regulates local lysosome availability in neurons. Somatic ER tubules control lysosome size and axonal translocation by promoting lysosome homo-fission. ER tubule – lysosome contacts are enriched in a pre-axonal region, where the kinesin-1-binding ER-protein P180 interacts with axonal MTs to promote kinesin-1-dependent lysosome translocation into the axon.

### Somatic ER tubule – lysosome contacts in axonal lysosome availability

Both the ER and lysosomes play essential roles in neuronal development and maintenance, and their distribution and organization must be tightly regulated to meet local demands. We have acquired a better understanding of the importance of MT-driven motor – organelle coupling for neuronal local availability of these two organelles, but only by studying each organelle in isolation (Farías et al., 2017; Farías et al., 2019). The ER and lysosomes are both distributed along the somatodendritic and axonal domains, and contacts between these two organelles have been visualized in unpolarized cells and neurons (Friedman et al., 2013; Wu et al. 2017). How neuronal organelle availability is regulated by local organelle organization and communication via organelle-organelle contacts is an outstanding question. Here, we found that a balance between ER tubules and ER cisternae is required for proper axonal distribution of lysosomes. The conversion between tubules and cisternae is regulated by ER tubule-shaping proteins such as RTNs and DP1, and the ER cisternae-shaping protein CLIMP63 (Voeltz et al., 2006; Shibata et al., 2010). We observed that knockdown of the ER-shaping proteins RTN4 and DP1 causes a reduced lysosome distribution in the axon, while knockdown of CLIMP63 increased axonal lysosome distribution. A similar phenotype has been observed for ER distribution in neurons, in which ER tubule disruption decreases axonal ER distribution, while ER cisternae disruption increase axonal distribution of ER tubules in the axon (Farías et al., 2019). This initially led us to speculate that axonal ER tubules contribute to the abundance of axonal lysosomes. However, axonal ER tubule repositioning into the soma did not affect the distribution or transport of lysosomes along the axon. On the contrary, we found that somatic ER tubule redistribution into the axon, caused impaired axonal lysosomes translocation. Interestingly, a previous EM study in brain tissue showed that ER – lysosome contacts mainly occur in the soma (Wu et al., 2017), a finding that we further confirmed using the split APEX assay to visualize contact sites. Together this suggests that the importance of somatic ER tubule organization in regulating axonal lysosome translocation is mediated by ER – lysosome contacts. Indeed, we found that ER tubule organization is required to form these contacts with lysosomes. ER – lysosome contacts could dynamically assemble and disassemble, as lysosomes were not dragged along by a sustained ER tubule relocation towards the axon. Several ER – organelle tethering proteins have been identified at contact sites, with the ER protein VAP playing a main role in ER tethering to multiple organelles as well as the plasma membrane (Wu et al., 2018). We analyzed the role of VAP in axonal lysosome distribution by knockdown experiments; however, we found only a modest reduction in axonal distribution after knockdown of both VAP-A and VAP-B (Figures S2A and S2B). Although several tether proteins pairs have been identified in ER – endosome or lysosome contact sites, their knockdown does not always result in contact loss (Rocha et al., 2009; Alpy et al., 2013; Eden et al., 2016; Fowler et al., 2019; Lee and Blackstone, 2020). This suggest that other tethering molecules may be involved or compensate for contact formation in neurons to regulate axonal lysosome translocation. Identifying neuron-specific tethering proteins involved in the formation and maintenance of ER – lysosome contact sites will be an important future research goal.

### ER tubules regulate lysosome size and motility

ER – endosome contacts increase as endosomes mature into a lysosome (Friedman et al., 2013). These contacts have been shown to promote endosome fission in non-neuronal cells (Rowland et al., 2014). Here, we have shown that ER tubule disruption causes enlarged and less motile mature lysosomes. Hundreds of fusion and fission events between mature and immature lysosomes were observed in the soma of control neurons in a period of 300 seconds, while ER tubule disruption caused a drastic reduction of around 80% in lysosome fission events. This indicates an important role for ER tubules in lysosome fission in order to maintain proper lysosome size and number to meet local demands in neurons.

A recent study has reported that knockdown of spastin, a MT-severing protein associated to ER tubules, also results in impaired endosome fission and enlarged lysosomes. Spastin and actin nucleators, such as the WASH complex component strumpellin, could generate the environment to promote lysosome constriction and fission at ER tubule – lysosome contact sites (Allison et al., 2017). In the same study, they also observed increased secretion of lysosomal enzymes into the extracellular space, suggesting impaired trafficking of enzymes into lysosomes (Allison et al., 2017). However, we have detected enzyme activity (active Cathepsin B and D) within enlarged lysosomes in live neurons and the presence of intraluminal vesicles in enlarged lysosomes by CLEM, indicating that these membranes are subject to lysosomal degradation. In our study, enlarged mature lysosomes were often less motile after ER tubule disruption, suggesting there may also be impaired coupling to the kinesin-1 motor. Besides its function in lysosome translocation, kinesin-1 was also shown to be involved in lysosome fission (Du et al., 2016). Consistent with this, we found that disruption of the kinesin-1-binding ER protein P180 caused a drastic reduction in lysosome motility and the enlargement of mature lysosomes, although they were smaller compared to ER tubule disruption. Since P180 knockdown drastically affected lysosome motility, the observed impairment in both lysosome fission and fusion could be explained by an initial defect in lysosome fission, as lysosomes often agglomerated and were unable to completely separate and translocate to fuse with another lysosome. It is possible that ER – lysosome contacts stabilize the parent lysosome to facilitate transfer of the kinesin-1 motor to the budding lysosome, which can then generate the forces to complete the fission and promote its subsequent translocation.

### ER – lysosome – MT interplay at a pre-axonal region in axonal organelle translocation

Interestingly, we observed a striking enrichment of contacts between the ER and lysosomes at a pre-axonal region. This region is featured by the landing of the kinesin-1 motor on stable MTs, where it is required for lysosome and ER tubule translocation into the axon (Farías et al., 2015; Farías et al., 2017; Farías et al., 2019). We previously found that the ER protein P180 is enriched in axonal ER tubules at a region preceding the axon initial segment, and it is involved in ER – MT co-stabilization (Farías et al., 2019). Here we show that, in this same region, P180 associates with stable MTs decorated by the kinesin-1 KIF5A-rigor mutant and that P180 is required for lysosome translocation into the axon. We observed an ER ring rearrangement around lysosomes at a pre-axonal region. ER rings around lysosomes have previously often been observed interacting with MTs, and they reduce diffusive motility of lysosomes (Friedman et al., 2013; Guo et al., 2018). In the absence of contacts with the ER, even when bound to MTs, lysosomes tend to undergo diffusive movement rather than directional transport along MTs (Guo et al., 2018). Guo et al proposed that ER – lysosome interactions may assist in stabilizing lysosomes prior their docking onto MTs via molecular motors (Guo et al., 2018). Consistent with a possible role of P180 in this process, we observed a drastic impairment in lysosome directional motility in P180 knockdown neurons. The cytoplasmic tail of P180 contains a MT-BD in a basic decapeptide repeat region, and a KIF5-BD in a coiled-coil (CC) region at the end of the C-terminal tail (Ogawa-Goto et al., 2007; Diefenbach et al., 2004). In neurons, expression of a P180ΔCC deletion construct containing the MT-BD, but not the KIF5-BD, promotes ER tubule – MT co-stabilization and distribution of P180 along the axon (Farías et al., 2019). In this study, we find that both the MT-BD and KIF5-BD are required for proper lysosome translocation into the axon. The role of P180 is likely downstream of ER tubule formation and ER tubule – lysosome contacts formation, as knockdown of P180 did not result in an evident reduction in contact formation. Concordantly with a previous finding of kinesin-1 mediating lysosome fission (Du et al., 2016), we observed that P180 disruption produces an agglomeration of less motile lysosomes mainly in a pre-axonal region, which were unable to separate from each other and translocate into the axon. We propose a multi-step model, in which the MT-BD of P180 locally stabilizes the interaction of ER tubule – lysosome contacts with MTs at a pre-axonal region. Then, the kinesin-1-BD of P180 facilitates kinesin-1 loading onto the budding lysosome, while part of the lysosome remains stabilized by the ER – MT interaction. Loading of kinesin-1 onto the budding lysosome at a pre-axonal region could locally facilitate the final step in lysosome fission and promote its subsequent translocation into the axon.

Other studies in cell lines have shown that protrudin, another kinesin-1-binding ER-protein promotes kinesin-1 loading onto lysosomes (Raiborg et al., 2015a). We find that protrudin is distributed in ER tubules wrapping lysosomes at a pre-axonal region; however knockdown of protrudin did not impair lysosome translocation into the axon. Because of the possible redundancy in tethering proteins, we cannot discard the involvement of protrudin in contact formation and motor loading onto lysosomes in neurons.

Together, our results support a model in which ER tubule – lysosome contacts interact with stable axonal MTs at a pre-axonal region to locally promote kinesin-1-powered lysosome fission and subsequent kinesin-1-mediated translocation into the axon. More broadly, our results suggest that organelle organization, inter-organelle communication and organelle transport are finely orchestrated to control local organelle availability in neurons. The fact that several ER-shaping proteins and contact tethering proteins are mutated in the neurodegenerative diseases hereditary spastic paraplegia and amyotrophic lateral sclerosis, highlight the importance of ER organization and inter-organelle communication in neuronal health (Fowler et al., 2019; Lee and Blackstone, 2020).

## ACKNOWLEDGMENTS

We thank Dr. Juan Bonifacino (NIH) for sharing the GFP-KIF5A-rigor and mCh-KIF5A-Strep constructs. We acknowledge C. de Heus and T. Veenendaal of the Cell Microscopy Center UMC Utrecht for their valuable assistance with CLEM experiments. We thank Dr. Lukas Kapitein for discussion during the project and Dr. Anna Akhmanova for critically reading the manuscript. This work was supported by the Netherlands Organization for Scientific Research (NWO) through a VIDI grant (016.Vidi.189.019) to G.G.F; Alzheimer Nederland (WE. 15045) to C.C.H; NWO through a ZonMW-TOP grant (91216006), Alzheimer Nederland (WE.03-2019-10), NWO Roadmap on Netherlands Electron Microscopy infrastructure NEMI (project 184.034.014) and the Deutsche Forschungs Gemeinschaft (DFG FOR2625) to JK.

## AUTHOR CONTRIBUTIONS

N.Ö. designed and performed experiments, analyzed data, and wrote the manuscript; M.K. designed and performed experiments, analyzed data, and wrote the manuscript; I.v.S. performed experiments related to enlarged lysosome after ER tubule disruption; A.v.H. performed experiments related to P180; N.L. designed and performed correlative light and electron microscopy experiments; J.K. discussed data, provided feedback and edited the manuscript; C.C.H. proposed experiments, discussed data and provided feedback and edited the manuscript; G.G.F. designed and performed experiments, analyzed data, supervised the research, coordinated the study, and wrote the manuscript.

## METHODS

### CONTACT FOR REAGENT AND RESOURCE SHARING

Further information and requests for resources and reagents should be directed to and will be fulfilled by the Lead Contact Ginny Farías.

### EXPERIMENTAL MODEL AND SUBJECT DETAILS

#### Animals

All experiments were approved by the DEC Dutch Animal Experiments Committee (Dier Experimenten Commissie), performed in line with institutional guidelines of University Utrecht, and conducted in agreement with Dutch law (Wet op de Dierproeven, 1996) and European regulations (Directive 2010/63/EU). Female pregnant Wistar rats were obtained from Janvier, and embryos (both genders) at E18 stage of development were used for primary cultures of hippocampal neurons. The animals, pregnant females and embryos have not been involved in previous procedures.

#### Primary neuronal cultures and transfection

The hippocampi from embryonic day 18 rat brains were dissected and dissociated in trypsin for 15 min and plated on coverslips coated with poly-L-lysine (37.5 μg/mL) and laminin (1.25 μg/mL) at a density of 100,000/well or 50,000/well (12-well plates) to prepare primary hippocampal neurons. Neurobasal medium (NB) supplemented with 1% B27 (GIBCO), 0.5 mM glutamine (GIBCO), 15.6 μM glutamate (Sigma), and 1% penicillin/streptomycin (GIBCO) was used to maintain the neurons incubated under controlled temperature and CO_2_ conditions (37°C, 5% CO_2_). Hippocampal neurons were transfected using Lipofectamine 2000 (Invitrogen). Briefly, DNA (0,05-2 μg/well) was mixed with 1.2 μL of Lipofectamine 2000 in 200 μL Opti-MEM, incubated for 20 min at room temperature, then added to neurons in NB and incubated for 1 hour at 37°C in 5% CO2. Next, neurons were washed with NB and transferred to their original medium at 37°C in 5% CO2 until fixation or imaging.

### METHOD DETAILS

#### DNA and shRNA Constructs

The following vectors were used: pEGFP(A206K)-N1 and pEGFP(A206K)-C1 (a gift from Dr. Jennifer Lippincott-Schwartz), pGW1-mCherry and pGW1-BFP (Kapitein et al., 2010) and pSuper (Brummelkamp et al., 2002). GFP-KIF5A-Rigor, LAMP1-GFP and mCherry-KIF5A-motor-Strep were a gift from Dr. Juan Bonifacino (Farías et al., 2015; Farías et al., 2017) and RFP-CLIMP63 was a gift from Dr. Tom Rapoport. RTN4A-GFP was provided by Dr. Gia Voeltz (Shibata et al., 2008; Addgene plasmid #61807). V5-GFP-P180 full length (Hung et al., 2017; Addgene #92150), TOM20-V5-FKBP-split-AP and Split-EX-HA-FRB-CB5 (Han et al., 2019; Addgene #120914 and #120915, respectively) were provided by Dr. Alice Ting. LAMP1-RFP was provided by Dr. Walther Mothes (Sherer et al., 2003; Addgene #1817). GFP-Rab7a and GFP-Rab11a were previously described (Hoogenraad et al., 2010).

For Strep/SBP heterodimerization system, the cloning of HA-KIFC1-MD-Strep and GFP- or mCh-SBP-RTN4A has been previously described (Farías et al., 2019). P180ΔMT-BD-GFP construct corresponds to a deletion construct lacking the entire P180 decapeptide repeat domain (containing the MT-BD), and it was previously described (named as P180-Δrepeat-GFP in Farías et al., 2019). We were unable to generate a deletion construct lacking only the MT-BD because of the nature of the repeated decapeptide sequence present in this domain of P180.

The plasmids generated in this study include:

For HA-KIF5A-Strep, the mCherry sequence from mCherry-KIF5A-motor-Strep (Farías et al., 2015) was removed by digestion with AgeI and BsrGI enzymes and replaced by a 3x HA sequence.

Primers used to anneal the 3x HA sequence were as follows:

5’ccggtgcccaccatgtacccatacgatgttcctgactatgcgggctatccctatgacgtcccggactatgcaggatcctatccatatgacgttccagattacgctggatccgt -3’ and
5’gtacacggatccagcgtaatctggaacgtcatatggataggatcctgcatagtccgggacgtcatagggatagcccgcatagtcaggaacatcgtatgggtacatggtgggca-3’

For P180-mCherry, full length P180 was PCR amplified from V5-GFP-P180 (Addgene #92150) and inserted in mCherry-N1 vector between XhoI and BamHI sites. A 3x(glycine-serine) linker was generated by addition to the cloning primers and was introduced between P180 and before the mCherry sequence to allow freedom of movement between domains.

For P180 deletion construct P180-ΔKIF5-BD-GFP, DNA sequences between nucleotides 1-3877 and 4236-4617 were PCR amplified from V5-GFP-P180 (Addgene #92150) and the two fragments were assembled and cloned into pEGFP(A206K)-N1 between XhoI and BamHI sites by GIBSON assembly. A 3x(glycine-serine) linker was introduced between fragments and before the GFP sequence. The primers used to generate P180ΔKIF5-BD-GFP construct were:

5’-agcgctaccggactcagatctcgagcaccatggatatttacgacactcaaaccttgggggttgtgg-3’ and
5’-ccgctgccgctacctgcggcgcccaccttggc-3’ for fragment 1,
5’-gggcgccgcaggtagcggcagcggtagcgagcaggaccccgttcagctg-3’ and
5’caccatggtggcgaccggtggatccgggctaccgctgccgctacccacgctggtgccctcctt-3’ for fragment 2.

For the Split APEX assay, we generated Split-AP-V5-protrudin and Split-EX-HA3x-Rab7a as follows: First, the GFP sequence in GFP-C1 vector was removed and replaced with Split-AP-V5 and Split-EX-HA3x sequence between AgeI and BglII sites. To generate Split-AP-V5-C1 vector, V5 and AP fragments were amplified from GFP-V5-P180 (Hung et al., 2017; Addgene # 92150) and TOM20-V5-FKBP-split-AP (Han et al., 2019; Addgene# 120914), respectively. A 3x(glycine-serine) linker was introduced before and after the V5 sequence. To generate Split-EX-HA3x-C1 vector, EX and HA fragments were amplified from Split-EX-HA-FRB-CB5 (Han et al., 2019; Addgene# 120915) and HA3x-KIF5A-Strep (generated in this study), respectively. A 3x(glycine-serine) linker was introduced between EX and HA fragments. Then, to generate Split-AP-V5-protrudin and Split-EX-HA-Rab7, human protrudin sequence was PCR amplified from IMAGE 4818199 (SourceBioScience) and Rab7a sequence was PCR amplified from GFP-Rab7 (Hoogenraad et al., 2010). Both Protrudin and Rab7 were cloned into Split-AP-V5-C1 and Split-Ex-ln-HA3x-C1 vectors, respectively, between XhoI and EcoR1 sites by GIBSON assembly. A 3x(glycine-serine) linker was introduced before the Rab7a sequence.

The following sequences for rat-shRNAs, inserted in pSuper vector, were used in this study: RTN4-shRNA (5’-gtccagatttctctaatta-3’), DP1-shRNA (5’-gacatataaagttccagaa-3’), P180-shRNAs (5’-tcagtgcaattgtctgtat-3’ and 5’-taaaccaaccaacacagcg-3’), KTN1-shRNA (5’-ggaccttctcaagaggtta-3’), and CLIMP63-shRNA (5’-tcaaccgtattagtgaagttctaca-3’) (Farías et al., 2019); VAPA-shRNA (5’-gcatgcagagtgctgtttc-3’) and VAPB-shRNA (5’-ggtgatggaagagtgc-3’) (Teuling et al., 2007; Lindhout et al., 2019). A previously described sequence for Protrudin-shRNA (5’-aagcttcttgatccgactggaag-3’; Shirane and Nakayama, 2006) was cloned into pSuper vector after oligo annealing.

#### Antibodies and reagents

The following primary antibodies were used in this study: rabbit anti-LAMTOR4 (Cell Signaling, clone D6A4V, Cat# 12284S, RRID: AB_2797870), mouse anti-EEA1 (BD Biosciences, Cat# 610456, RRID: AB_397829), mouse anti-P62 (Abcam, Cat# 56416, RRID:AB_945626), mouse anti-V5 (Thermo Fisher Scientific Cat# R960-25, RRID:AB_2556564), rat anti-HA (Roche Cat# 11867423001, RRID:AB_390918), mouse anti-Pan-Neurofascin external (clone A12/18; UC Davis/NIH NeuroMab, Cat# 75-172, RRID: AB_2282826), and rabbit anti-TRIM46 (van Beuningen et al., 2015)

The following secondary antibodies were used in this study: Streptavidin, Alexa Fluor 555 conjugate (Thermo Fisher Scientific Cat# s21381, RRID: AB_2307336), Streptavidin, Alexa Fluor 568 conjugate (Thermo Fisher Scientific Cat# S-11226, RRID:AB_2315774), donkey anti-mouse Alexa488 (Molecular Probes, Cat# A21202, RRID: AB_141607), donkey anti-mouse Alexa555 (Molecular Probes, Cat# A31570, RRID: AB_2536180), donkey anti-mouse Alexa647 (Molecular Probes, Cat#A31571, RRID: AB_162542), donkey anti-rabbit Alexa488 (Molecular Probes, Cat# A21206, RRID: AB_141708), donkey anti-rabbit Alexa555 (Molecular Probes, Cat# A31572, RRID: AB_162543), donkey anti-rabbit Alexa647 (Molecular Probes, Cat# A31573, RRID: AB_2536183), goat anti-mouse Alexa405 (Molecular Probes, Cat# A31553, RRID: AB_221604), goat anti-rabbit Alexa405 (Molecular Probes, Cat# A31556; RRID: AB_221605), goat anti-rat Alexa 488 (Thermo Fisher Scientific Cat# A-11006, RRID:AB_2534074), goat anti-rat Alexa 568 (Thermo Fisher Scientific Cat# A-11077, RRID:AB_2534121).

Other reagents used in this study were NeutrAvidin (Thermo Fisher Scientific, Cat# 31000), Lipofectamine 2000 (Invitrogen, Cat#1639722), SiR-lysosome kit (Spirochrome, Cat# SC012), Magic Red (ImmunoChemistry Technologies, Cat# 937); antibody labeling kit Mix-n-Stain CF640R (Biotium); heme (Sigma-Aldrich, Cat#51280); biotin-phenol (Iris Biotech, Cat#LS.3500); H2O2 (Sigma-Aldrich, Cat#H1009)

#### Immunofluorescence staining and imaging

Neurons were incubated at RT with pre-warmed 4% paraformaldehyde plus 4% sucrose in PBS for 20 min for fixation. Then, cells were permeabilized with 0.2% Triton X-100 in PBS supplemented with calcium and magnesium (PBS-CM) for 15 min, followed by blocking with 0.2% porcine gelatin in PBS-CM for 30 min at 37°C. Next, neurons were incubated with primary antibodies and then with secondary antibodies for 30 min at 37°C each. After incubation with primary and secondaries antibodies, the cells were washed with PBS-CM 3 times for 5 min each. Coverslips were mounted in Fluoromount-G Mounting Medium (ThermoFisher Scientific) and imaged by using a confocal laser-scanning microscope (LSM700, Zeiss) equipped with Plan-Apochromat 63x NA 1.40 oil DIC and EC Plan-Neofluar 40x NA1.30 Oil DIC objectives.

#### Labelling mature lysosomes

Prior to live-cell imaging, DIV7 hippocampal neurons were incubated with SirLyso (1000 nM in NB; Spirochrome) to detect cathepsin D activity, or Magic-Red (1:250 dilution in NB from recommended stock reconstruction; ImmunoChemistry Technologies) to detect cathepsin B activity. Both probes were incubated for 30 minutes under controlled temperature and CO_2_ conditions (37°C, 5% CO_2_). After washing twice with NB, cells were supplemented with their original medium and immediately imaged.

#### Correlative Light and Electron Microscopy

For correlation of FM and 3D-EM of neurons, FM imaging was performed prior to sample preparation for EM. Neurons were cultured on carbon-coated, gridded coverslips. DIV7 neurons incubated with SirLyso were rinsed and fixed with 4% paraformaldehyde plus 4% sucrose in 0.1M PB for 120 min. Coverslips were imaged in fixative solution by using a confocal laser-scanning microscope (LSM700, Zeiss) equipped with Plan-Apochromat 63x NA 1.40 oil DIC objective. The position of cells relative to the pattern etched in the coverslip was registered using polarized light. After fluorescent imaging, neurons on coverslips were post-fixed with 1% OsO4 with 1.5% K4Fe(II)(CN)6 in 0.1M PB for 1 hour on ice, followed by washing steps in ddH2O. Cells were stained with 2% uranyl acetate in ddH2O at room temperature, followed by further washing steps with ddH2O. Finally, samples were subjected to a graded ethanol series for dehydration. After dehydration, samples were flat embedded in Epon resin (ratio: 12g Glycid Ether 100, 8g dodecenylsuccinic anhydride, 5.5g methylnadic anhydride, 560 μL N-benzyldimethylamine). After Epon polymerization, the resin blocks were removed from the coverslips and prepared for EM as reported before (Fermie et al., 2018) with slight modifications. Regions of interest selected based on fluorescent imaging (LAMP1-GFP and SirLyso) were cut out using a clean razor blade, and glued to empty Epon sample stubs, with the basal side of the cells facing outwards. The resin embedded neurons were then mounted on aluminum SEM stubs using carbon adhesive, and the sides of the block were covered with conductive carbon paint. Samples were imaged using a Scios Dualbeam FIB-SEM (Thermo Fischer Scientific) under high vacuum conditions. A 500 nm thick Pt layer was deposited over the ROI using the FIB (30kV, 1 nA). Then the trenches around the selected ROI were milled, and the imaging surface was polished. Automated serial imaging was performed using Slice&View v3 (Thermo Fischer Scientific), at low acceleration voltages (2 kV) using 5 nm pixel size, dwell time 5μs at a slice thickness of 5 nm providing isotropic pixels in 3D. Backscattered electrons were collected using the in-lens backscatter detector operating in ‘Optitilt’ mode. Images were saved as separate 8-bit TIFF files.

The resulting images were imported in Fiji (Fiji is just ImageJ) to generate 3D volumes as a single stack and aligned using Fiji Plugin SIFT. Aligned XZ stacks were reconstructed as XY stacks (FM imaging plane) and saved as a single TIFF and converted to MP4 (Videos S7 and S8). Aligned and reconstructed slices were manually registered over fluorescent images. For correlation of FM and EM data, the best matching XY plane from the reconstructed stack of the ROI was overlayed with FM data using Photoshop. Multiple corresponding spots (e.g. lysosomes) on images were selected and overlay of FM and EM data was generated by linear scaling and transformation steps were followed. Only linear transformation options were used to achieve the overlays shown in the Figure 3K. The measurement of lysosome diameter was also performed in Fiji, using the line segment tool.

#### Live-cell imaging

For live-cell imaging experiments, an inverted microscope Nikon Eclipse Ti-E (Nikon), equipped with a Plan Apo VC 100x NA 1.40 oil and a Plan Apo VC 60x NA 1.40 oil objective (Nikon), a Yokogawa CSU-X1-A1 spinning disk confocal unit (Roper Scientific), a Photometrics Evolve 512 EMCCD camera (Roper Scientific) or Photometrics Prime BSI camera, and an incubation chamber (Tokai Hit) mounted on a motorized XYZ stage (Applied Scientific Instrumentation) was used. MetaMorph (Molecular Devices) software was installed for controlling all devices. Coverslips mounted in a metal ring and supplemented in the original medium from neurons were imaged in an incubation chamber that maintains optimal temperature and CO2 (37°C and 5% CO2). To visualize proteins with a specific fluorescent tag for single-color acquisition, a laser channel was exposed for 100-200 ms while for dual-color acquisition, different laser channels were exposed for 100-200 ms sequentially. Neurons were imaged every 1 sec for 300 sec. To identify the axon, neurons were incubated with a CF640R-conjugated antibody against the AIS protein neurofascin (NF-640R; Farías et al., 2016) for 30 min before live-cell imaging. Total time and intervals of imaging acquisition for each experiment are depicted in each legend for Figure and/or legend for Video.

#### Streptavidin/SBP heterodimerization system assay

Controlled coupling between MT-driven motor proteins and a specific cargo such as vesicles, lysosomes, and ER tubules, using the Strep/SBP heterodimerization system, has been previously described (Farías et al., 2015; Farías et al., 2017; Farías et al., 2019). Shortly, neurons were transfected at DIV5 with Strep-KIFC1-MD-HA plus GFP-SBP-RTN4A to pull axonal ER tubules to soma (Figures 2B and 2D; Videos S2 and S4) or HA-KIF5A-Strep plus GFP-SBP-RTN4A to pull ER tubules from the soma into axon (Figures 2C, 2D and 3C; Videos S2, S4 and S5). Strep-SBP uncoupling was prevented by adding NeutrAvidin (0.3mg/mL) to the cell medium after 1h of transfection (Farías et al., 2016).

#### Split APEX assay

Neurons were transfected at DIV4 with AP-V5-Protrudin and EX-3xHA-Rab7 constructs. At DIV7, a final concentration of 6uM heme (Sigma-Aldrich) was added to the medium and after 60 min neurons were washed once with NB and 500μM biotin-phenol (Iris Biotech) in NB with supplements was added to the neurons for 30 min. Then, proximity labeling was initiated by adding H2O2 (Sigma-Aldrich) to a final concentration of 1mM for 1 minute after which the labeling reaction was stopped by removing the medium and washing once with quenching buffer (5 mM Trolox (Sigma-Aldrich) and 10 mM sodium ascorbate (Sigma-Aldrich) in HBSS) containing 10 mM sodium azide (Merck) and twice with quenching buffer without sodium azide for 3-5 min each. Neurons were subsequently fixed and stained as described above.

#### Image analysis and quantification

Images were recorded and analyzed from 3-5 independent experiments. No specific strategy for randomization and/or stratification was employed. Data was analyzed for at least two people in a blind fashion.

##### Fluorescence line intensity plots

The co-distribution of different markers was analyzed using ImageJ. Plot profiles were generated from lines traced along lysosomes (Figures 3D, 5B, S1A, S1B, S1C, S1D and S1E), or segmented line traced from a somatic pre-axonal region to the proximal axon (Figure 6E). The length of traced line is indicated in each intensity plot.

##### Polarity index of lysosomal markers

Quantification of polarity index was performed using ImageJ, as previously described (Farías et al., 2019). Shortly, segmented lines were drawn along three dendrites and one portion of the axon of ~200 μm (excluded the axon initial segment) in each image. Mean intensities in these areas were measured by ImageJ. After averaging the mean intensities from the three dendrites, following formula was applied to calculate the polarity index:*PI* = (*Id* − *Ia*)/(*Id* + *Ia*): in which Id is the average intensity of the three dendrites and Ia is the intensity of axon. PI<0 indicates axonal distribution, PI>0 indicates dendritic distribution and PI=0 stands for non-polarized distribution where *Id*=*Ia* (Figures 1B, 1D and S2B).

##### Kymograph analysis

Kymographs from live cell images were made using Image J as previously described (Farías et al., 2016). Shortly, segmented lines were drawn along a 30-μm segment of the axon from the most distal part of the axon initial segment as indicated in schematic in Figure 1F. Then regions were strengthened and re-sliced followed by z-projection to obtain kymograph. Anterograde movements were oriented in all kymographs from left to right. Time of recording and length of segments are indicated in each kymograph (Figures 1E, 2D, 6A and 6H). Number of events for antero- and retrograde lysosome movement as well as for stationary lysosomes were obtained from kymographs from many cells (Figures 1F, 2E, 6B and 6I).

##### Quantification of number and size of lysosomes

The number of LAMP1-positive lysosomes and LAMP1/SirLyso-positive mature lysosomes as well as the size of mature lysosomes was analyzed using ImageJ. The number of lysosomes were counted manually from the first frame of live soma images of 7-8 different neurons per condition by three independent observers. In total, we counted 845 LAMP1-positive and 588 SirLyso-positive lysosomes from 8 different control neurons, 411 LAMP1-positive and 195 SirLyso-positive lysosomes from 8 different RTN4/DP1 KD neurons and 625 LAMP1-positive and 421 SirLyso-positive lysosomes from 7 different P180 KD neurons. We plotted the average per neuron in Figures 3E and S2G). We calculated the ratio of mature/immature lysosomes by dividing the total amount of SirLyso-positive lysosome per soma by the total amount of LAMP1-positive lysosomes per soma (Figures 3F and S2H). To measure lysosome size, straight lines were traced along the diameter of spherical lysosomes from images of the soma from live neurons. The largest lysosome per soma was measured and averaged per condition (Figure 3I). We considered lysosomes with a size bigger than 1 μm as enlarged lysosomes (de Araujo et al., 2020). The total number of enlarged LAMP1/SirLyso-positive lysosomes were counted manually from the first frame of live soma images. We plotted the average per neuron in Figure 3G. The percentage of enlarged lysosomes was calculated by dividing the number of enlarged lysosomes (>1μm) by the total number of LAMP1/SirLyso-positive lysosomes per soma (Figure 3H). We used the same control neurons for comparisons with RTN4/DP1 KD neurons or P180 KD neurons in all analyses as the experiments were performed together.

##### Quantification of fusion and fission events

Homo-fusion and homo-fission events were analyzed using ImageJ. Merging of two LAMP1/SirLyso-positive mature lysosomes or one LAMP1/SirLyso-positive mature lysosome and one LAMP1-positive immature lysosomes were considered as fusion events while splitting of two LAMP1/SirLyso-positive mature lysosomes or splitting of LAMP1-positive lysosomes from LAMP1/SirLyso-positive mature lysosomes were considered as fission events. The number of fusion and fission events on all LAMP1/SirLyso-positive lysosomes (±588 in control, ±195 in RTN4/DP1 KD and ±421 in P180 KD) from the live soma images of 6-8 neurons per condition were counted manually for 301 frames (1 frame/sec) by three independent observers. The counts were averaged and plotted per soma (Figures 4K, 4L, 6K and S2C) or per lysosome by dividing the number of fusion or fission events by the total number of LAMP1/SirLyso-positive lysosomes per soma (Figures 4N, 4O, S2D and S2E). The fusion/fission ratio was calculated by dividing the total number of fusion events per soma by the total number of fission events per soma (Figures 4M and S2F). We used the same control neurons for comparisons with RTN4/DP1 KD neurons or P180 KD neurons in all analyses. as the experiments were performed together.

##### Quantification of immunofluorescence intensity for streptavidin

All images were taken with the same settings for light and exposure and with parameters adjusted so that the pixel intensities were below saturation. Quantification of the intensity of streptavidin signal was performed using ImageJ. z-projections of each image were generated using the average intensity and a ROI was manually drawn around the neuronal soma. Mean intensities from 16-bit images for one channel corresponding to streptavidin signal in the selected area was measured using ImageJ. Intensities were averaged over multiple cells and normalized to the average intensity in control cells (Figures 5G and 6J).

## STATISTICAL ANALYSIS

Data processing and statistical analysis were performed using Excel and GraphPad Prism (GraphPad Software). Unpaired and paired t-test, Kruskal-Wallis test followed by a Dunn’s multiple comparison test, Mann-Whitney U, one-way ANOVA test followed by a Tukey’s multiple comparisons test were performed for statistical analysis and are indicated in Figure legends. Significance as determined as followings: ns-not significant, *p<0.05 **p<0.01 and ***p<0.001. The assumption of data normality was checked using D’Agostino-Pearson omnibus test.

## LEGEND FOR SUPPLEMENTAL VIDEOS

**Video S1 (related to Figure 1). ER tubule disruption impairs lysosome translocation into the axon.** Transport of LAMP1-positive lysosomes in proximal axons of DIV7 neurons co-transfected with LAMP1-GFP, fill and control pSuper (top) or shRNAs targeting RTN4 plus DP1 (bottom). Endogenous NF mark the axon initial segment. Neurons were recorded every 1 sec for 300 sec. Orange dashed lines in left panels indicate region of the straightened proximal axon showed in right panels.

**Video S2 (related to Figure 2). Somatic ER tubule repositioning into the axon impairs axonal lysosome translocation**. Transport of LAMP1-positive lysosomes in proximal axons of DIV7 neurons transfected with only LAMP1 and SBP-RTN4 plus fill as a control (left panel), or together with Strep-KIFC1 (middle panel) to pull ER tubules from axon to soma, or KIF5A-Strep (right panel) to pull ER tubules from soma to axon. Neurons were imaged every 1 sec for 300 sec.

**Video S3 (related to Figure 3). Enlarged and less motile lysosomes in the soma of neurons after ER tubule disruption**. Lysosome motility in the soma of DIV7 neurons co-transfected with LAMP1-GFP, fill and control pSuper (top) or shRNAs targeting RTN4 plus DP1 (bottom). Neuron was recorder every 1 sec for 300 sec.

**Video S4 (related to Figure 3). Somatic ER tubule repositioning into the axon impairs lysosome size and motility**. Dynamics of LAMP1-positive lysosomes in the soma of DIV7 neurons transfected with only LAMP1 and SBP-RTN4A plus fill as a control (left) or together with Strep-KIFC1 (middle) or KIF5A-Strep (right). Neurons were recorded every 1 sec for 300 sec. Blue arrowhead point to the proximal axon.

**Video S5 (related to Figure 3). Somatic ER tubule disruption causes reduced motility of enlarged lysosomes in a pre-axonal region, thereby impairing axonal translocation**. Lysosome motility in neurons co-transfected with LAMP1, fill and control pSuper or shRNAs targeting RTN4 plus DP1. Neurons were recorded every 1 sec for 300 sec.

**Video S6 (related to Figures 3 and 4). Motile mature and immature lysosomes in the soma of control neurons and enlarged and less motile mature lysosomes in the soma after ER-tubule disruption**. Neurons were recorder every 1 sec for 300 sec.

**Video S7 (related to Figure 3). FIB-SEM slices in XZ orientation of part of the neurons shown in Figure 3 J-O**. The dataset shown is 10,76 μm in width.

**Video S8 (related to Figure 3). FIB-SEM slices in XY orientation of part of the neurons shown in Figure 3J-O, showing the morphology of the indicated lysosomes**. Images were captured at a lateral pixel size of 5 nm and 5 nm slice thickness, using the in-lens backscattered electron detector with inverted contrast. Horizontal width of the images in the stack is 6750nm.

**Video S9 (related to Figure 4). Lysosome fusion and fission events and impaired fission after ER tubule disruption**. Movies of fusion and fission events from still images shown in Figure 4.

**Video S10 (related to Figure 6). Knockdown of P180 leads to the accumulation of enlarged lysosomes at a pre-axonal region, thereby impairing axonal lysosome translocation**. Lysosome motility from still image shown in Figure 6C. Neuron was recorder every 1 sec for 300 sec. Axon initial segment (AIS) labelled with NF is shown in green dashed box. Enlarged and less motile lysosomes in a pre-axonal region is shown in orange dashed box.

**Video S11 (related to Figure 6). Enlarged mature lysosomes show reduced motility and accumulate in a pre-axonal region after P180 knockdown**. Motility of lysosomes in the soma and pre-axonal region of neurons after P180 knockdown. Neurons were recorder every 1 sec for 300 sec. Compare with control neurons shown in Video S6.

