## Supplemental Figures for "ER – lysosome contacts at a pre-axonal region regulate axonal lysosome availability"

### Figure S1

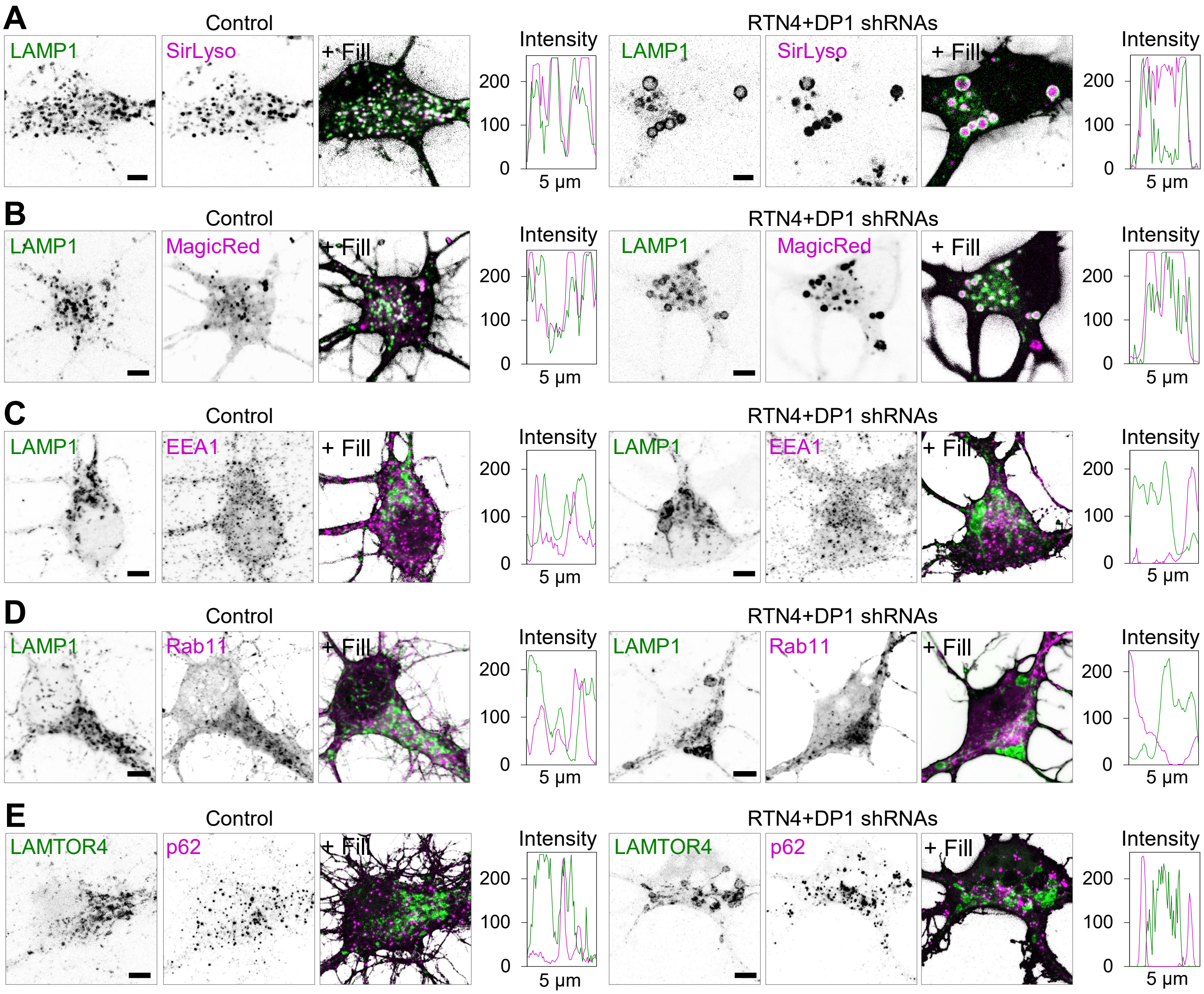
A
Control
LAMP1
SirLyso
+ Fill
Intensity
200
100
0
5 µm
RTN4+DP1 shRNAs
B
MagicRed
C
EEA1
D
Rab11
E
LAMTOR4
p62

**Figure S1 (related to Figure 3). Early endosome, recycling endosome and autophagosome markers are not particularly enriched in enlarged mature lysosomes after ER tubule disruption**

**(A-B)** Representative still images of the soma of DIV7 neurons transfected at DIV3 with a control pSuper plasmid (left panels) or pSuper plasmids containing shRNAs targeting RTN4 plus DP1 (right panels) together with LAMP1-GFP and a fill, and labelled live for active cathepsin-D and B (magenta) with SirLyso (A) and MagicRed (B), respectively. LAMP1 in green. Intensity profile line of a region of the soma for control and RTN4 plus DP1 knockdown neurons, on the right of each merged image.

**(C-E)** Representative images of lysosomes distributed in the soma of DIV7 neurons transfected at DIV3 with fill and a control pSuper plasmid (left panels) or pSuper plasmids containing shRNAs targeting RTN4 plus DP1 (right panels) (E), together with LAMP1-GFP (green) (C), or LAMP1-RFP (green) plus GFP-Rab11 (magenta) (D). Neurons were stained for EEA1 (early endosome marker, magenta) (C), or LAMTOR4 (lysosome marker, green) and p62 (autophagosome marker, magenta) (E). Intensity profile lines for different markers on the right of each merged image.

Scale bars represent 5 μm in (A), (B), (C), (D) and (E).

### Figure S2

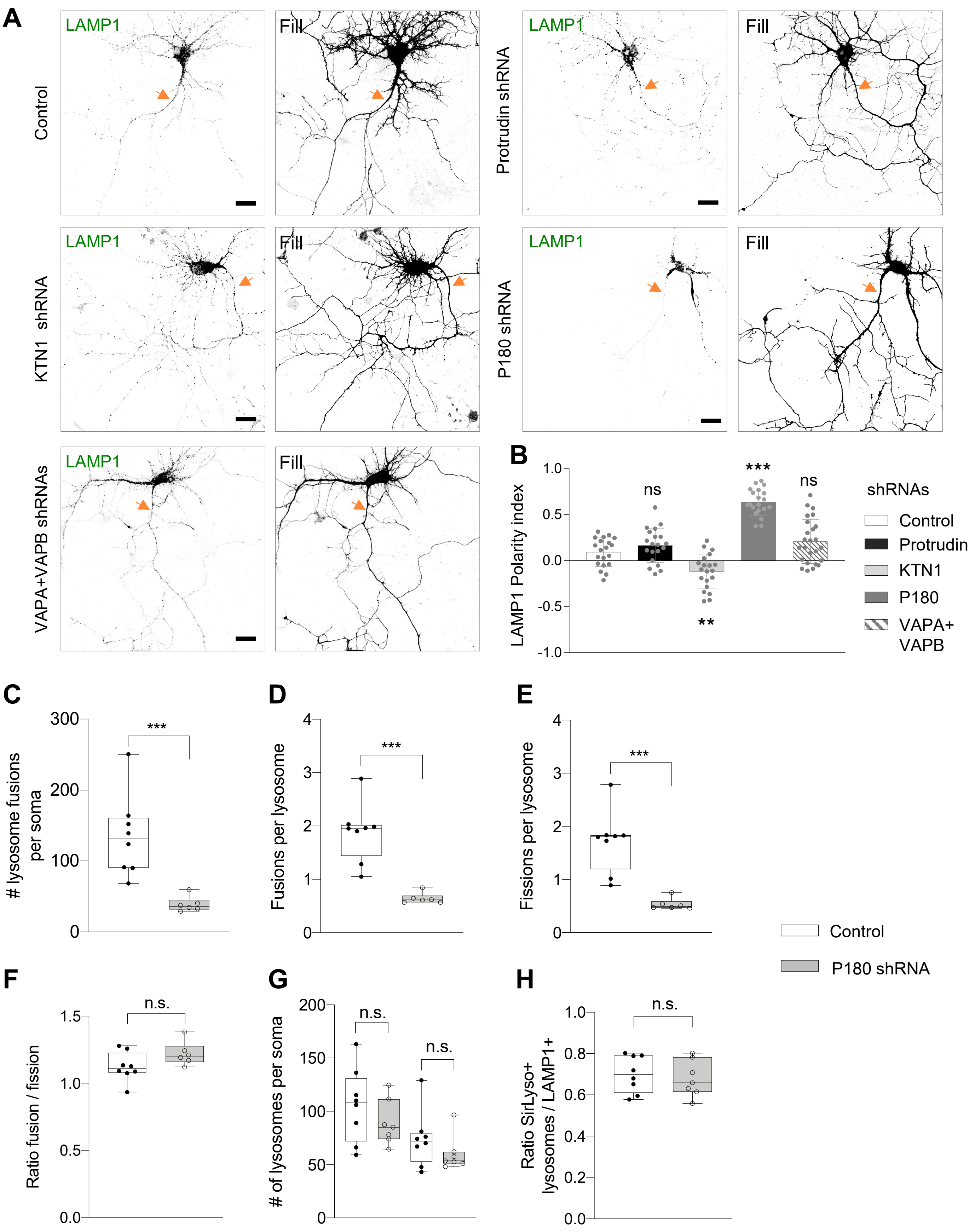
A
Control
LAMP1
Fill
Protrudin shRNA
KTN1 shRNA
P180 shRNA
VAPA+VAPB shRNAs
B
LAMP1 Polarity index
1.0
0.5
0.0
-0.5
-1.0
ns
**
***
ns
shRNAs
Control
Protrudin
KTN1
P180
VAPA+ VAPB
C
### lysosome fusions per soma
300
200
100
0
***
D
Fusions per lysosome
4
3
2
1
0
***
E
Fissions per lysosome
***
Control
P180 shRNA
F
Ratio fusion / fission
1.5
1.0
0.5
0.0
n.s.
G
### of lysosomes per soma
200
150
100
50
0
n.s.
n.s.
H
Ratio SirLyso+ lysosomes / LAMP1+
1.0
0.8
0.6
0.4
0.2
0.0
n.s.

**Figure S2 (related to Figure 6). Knockdown of the kinesin-1-binding ER-protein P180 impairs axonal lysosome distribution by reducing somatic lysosome motility, fusion and fission**

**(A-B)** Representative images of lysosome distribution in DIV7 neurons transfected with fill, LAMP1-GFP and pSuper plasmid or pSuper plasmids containing shRNA sequences targeting three kinesin-1-binding ER-proteins (protrudin, KTN1 and P180) as well as the tethering proteins VAPA and VAPB. Quantification of LAMP1 polarity indices in (B).

**(C-H)** Parameters indicated in each graph were quantified from neurons transfected with fill and a control pSuper plasmid or pSuper plasmids containing shRNAs targeting P180 together with LAMP1-GFP and stained with SirLyso on DIV7 prior to imaging for 300 s (1 frame/s). Control, white bars; P180 knockdown, gray bars. See also Video S9.

Scale bars in images represent 5 μm and orange arrows point to the proximal axon in (A). Boxplots show the mean and individual datapoints each represent a neuron; ns-not significant, $^{**}p<0.01$ and $^{***}p<0.001$ comparing conditions to control (ANOVA followed by a Dunn's multiple comparison test in (B), and (Mann-Whitney U) in (C), (D), (E), (F), (G) and (H).

### Figure S3

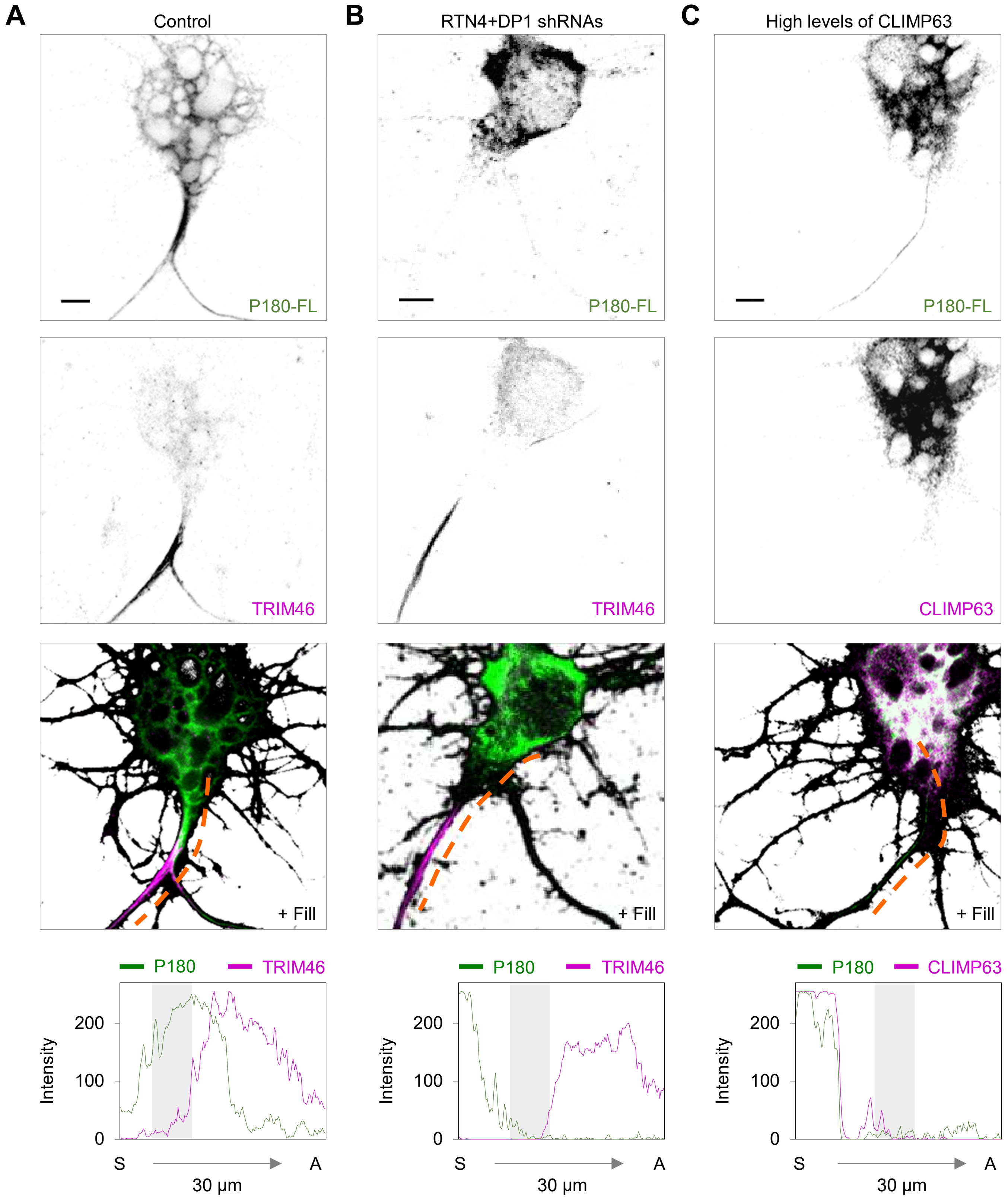

A
Control
B
RTN4+DP1 shRNAs
C
High levels of CLIMP63
P180-FL
P180-FL
P180-FL
TRIM46
TRIM46
CLIMP63
+ Fill
+ Fill
+ Fill
P180
TRIM46
P180
TRIM46
P180
CLIMP63
Intensity
200
100
0
S
A
30 µm

**Figure S3 (related to Figure 6) P180 requires ER tubule formation for its distribution in a pre-axonal region**

**(A-C)** Representative images of P180 distribution in the soma and pre-axonal region of DIV7 neurons transfected with fill and P180-GFP (green in merges) together with pSuper plasmid (A) or pSuper plasmids containing shRNAs targeting RTN4 plus DP1 (B) or high expression of the cisternae-shaping protein CLIMP63-RFP (magenta) (C) Axon initial segment was labelled with an antibody against TRIM46 (magenta in merges) in (A) and (B). Intensity profile lines from the soma to the proximal axon, indicated with orange lines in merged images, are shown in the bottom. Scale bars represent 5 μm.
